# Hypoxic Stress Induces LARP1-Driven Stress Granules That Protect mRNAs from Decay to Sustain Mitotic Cell Cycle Progression

**DOI:** 10.64898/2026.11.05.764444

**Authors:** Tiffanie Lim, Janani Ramanathan, Hongwei Gao, Erica Xu Songci, Tan Tuan Zea, Anthony Khong

## Abstract

Hypoxia is a defining feature of solid tumours that elicits extensive transcriptional and post-transcriptional adaptations. Stress granules (SGs) are frequently observed under hypoxia, yet how they assemble and whether they contribute to hypoxic gene regulation remain poorly understood. Here, we show that hypoxia induces SGs through an assembly mechanism distinct from that of classical acute stresses. Hypoxia SG formation occurs independently of the integrated stress response and instead requires the HIF1α–REDD1–mTORC1–4EBP1 axis. Remarkably, hypoxia SGs assemble despite robust ongoing translation, suggesting that additional mechanisms lower the threshold for RNA condensation. We identify LARP1 as a potential determinant of this process, where it promotes G3BP-dependent SG assembly specifically during hypoxia. Despite their distinct assembly mechanism, hypoxia and arsenite SGs recruit broadly similar RNA populations, with selective differences validated by single-molecule FISH, suggesting that conserved principles govern RNA partitioning across different SG states. Functionally, sequestration into hypoxia SGs correlates with enhance protein expression whereby a subset of mRNAs are protected from decay. Transcriptomic analyses further reveal that this protection sustains a broader gene-expression program enriched for MCM complex and mitotic cell-cycle regulators. Accordingly, disruption of hypoxia SGs reduces mitotic progression and proliferative competence during prolonged hypoxia. Together, our findings establish LARP1-mediated condensation as a mechanism that enables SG assembly despite ongoing translation and reveal that SG sequestration preserves selected mRNAs from decay to sustain the gene-expression programs required for long-term adaptation to hypoxia.

## INTRODUCTION

Hypoxia is a defining feature of solid tumours that profoundly remodels gene expression to promote cellular adaptation, tumour evolution, therapeutic resistance, and immune evasion^1^. While transcriptional responses mediated by the pVHL–HIF pathway are well established^2^, hypoxia also extensively alters mRNA processing, translation, and stability^3^, yet the mechanisms coordinating this post-transcriptional response remains an active area of study. One prominent post-transcriptional response to hypoxia is the assembly of stress granules (SGs)^4–6^, RNA–protein condensates that form in hypoxic cells and within hypoxic regions of solid tumours^4^ with unclear functions.

Classically, SGs form when cells are not engaged in translation. Specifically, when cells encounter a variety of stress conditions that elicit a strong integrated stress response (ISR) pathway, ribosomes are released from mRNAs, and these non-translating mRNPs reach a critical concentration threshold thereby promoting phase-separation from the rest of the cytosol in an energetically favourable manner^7–9^. Key proteins involved in this process includes G3BP1/2 which is thought to aid in LLPS mediated by its multivalency^10–12^, and also thought to promote RNA-RNA interactions as kinetic condenser^13^.

Despite well-understood model of SG assembly, the functions of SGs remain unclear and an active area of research. One model, the RNA triage model, purports that sequestration of RNA into SGs alters their fate by altering stability or translation^14^. However, most historical studies suggest SGs do not influence RNA fate^15–18^. But recent articles suggest SGs may actually influence RNA fate but the mechanisms involved is not completely well understood respectively^19,20^.

Herein, we address how hypoxia SGs assemble and what are their functions in SG-enriched RNAs in order to elucidate the biology of a granule present in tumour microenvironment, and to understand how SGs control post-transcriptional gene regulation fundamentally and in the hypoxia context. Our data indicate hypoxia SGs assemble independent of the ISR pathway but is dependent instead on HIF1α–REDD1–mTORC1– 4EBP1 axis. Surprising, unlike ISR-induced stresses, SGs can form in the presence of robust translational activity. Moreover, we discovered an additional RNA-BP, LARP1, may be involved in promote hypoxia SG condensation thereby potentially explaining how these granules can form with minimal translational shutoff. In addition, we learned that the RNAs sequestered to hypoxia SGs is protected from degradation. And these this protection facilitates mitotic cell cycle progression and MCM complex expression, thereby promoting long-term adaptation to hypoxia enabling proliferative competence.

## MATERIALS AND METHODS

### Cell culture

U-2 OS and HEK293T cells were cultured in T-75 flasks (Corning, 430641U) in high-glucose Dulbecco’s Modified Eagle Medium (DMEM; Thermo Fisher Scientific, 11965092) supplemented with 10% fetal bovine serum (FBS; Gibco, 16000-044) and 1% penicillin– streptomycin (Thermo Fisher Scientific, 15140122). Cells were maintained at 37°C in a humidified incubator containing 5% CO₂. For routine passaging, cells were dissociated using 0.5% trypsin–EDTA without phenol red (Thermo Fisher Scientific, 15400054), collected by centrifugation, and resuspended in fresh complete medium.

### Lentiviral generation and transduction

HEK293T cells were seeded at a density of 5 × 10⁵ cells in T-25 flasks (Corning, 430639) one day before transfection. Cells were transfected with the indicated transfer plasmids encoding GFP–G3BP1, Halo–USP10p^1–40^, Halo–USP10p^1–40(F10A)^, or short hairpin RNAs (shRNAs) targeting 4E-BP1 (TRCN0000335449 and TRCN0000075610; Sigma-Aldrich), together with the appropriate lentiviral packaging plasmids. Lentiviral plasmids pRP2985, pRP2986, pRP2987, and pRP2988 were kindly provided by the Parker laboratory. A non-targeting shRNA construct (Sigma-Aldrich, SHC016) was used as a control. Transfections were performed using Lipofectamine 2000 (Thermo Fisher Scientific, 11668019) and Opti-MEM (Thermo Fisher Scientific, 31985070) according to the manufacturer’s instructions.

Lentivirus-containing supernatants were collected 72 h after transfection and filtered through a 0.45 μm filter to remove cellular debris. U-2 OS cells were transduced with the harvested lentiviral supernatants in the presence of 1 μg/mL polybrene (Santa Cruz Biotechnology, sc-134220). The viral supernatant was replaced with fresh complete medium 1 h after transduction, and cells were cultured for an additional 3 days.

To establish cell populations stably expressing GFP–G3BP1, Halo–USP10p^1–40^, or Halo–USP10p^1–40(F10A)^, transduced cells were enriched by fluorescence-activated cell sorting (FACS). For Halo-tagged constructs, cells were incubated with 1× HaloTag ligand (Promega, GA1111) for 15 min at 37°C. The medium was subsequently replaced to remove unbound ligand, and cells were incubated for an additional 30 min before sorting. GFP–G3BP1-expressing cells were sorted based on intrinsic GFP fluorescence. Cell sorting was performed using a BD FACSAria III cell sorter (BD Biosciences) at the Flow Cytometry Facility, Cancer Science Institute of Singapore (CSI), National University of Singapore.

### siRNA transfection

For transient depletion of 4E-BP1, HIF-1α, REDD1, and LARP1, U-2 OS cells were seeded in six-well plates and transfected with the indicated small interfering RNAs (siRNAs) using Lipofectamine RNAiMAX (Thermo Fisher Scientific, 13778075) according to the manufacturer’s instructions. Non-targeting siRNAs were used as controls. Cells were subjected to the indicated stress treatments 48 h after transfection. The shRNA and siRNA reagents used in this study are listed in Tables 1 and 2, respectively.

**Table 1.**
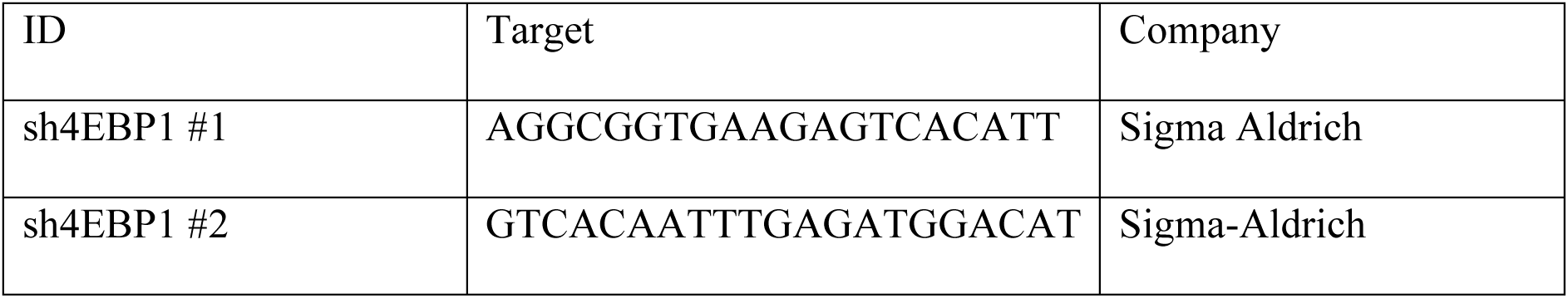
shRNA constructs used for stable gene knockdown.

**Table 2:** siRNAs used for transient knockdown.

| ID | Company | Catalogue No. |
|---|---|---|
| DsiHIF1α #1 | hs.Ri.HIF1A.13.1 | IDT |
| DsiHIF1α #2 | hs.Ri.HIF1A.13.2 | IDT |
| DsiLARP1 #1 | hs.Ri.LARP1.13.1 | IDT |
| DsiLARP1 #2 | hs.Ri.LARP1.13.2 | IDT |

### Stress treatments

U-2 OS cells were subjected to the indicated stress treatments for the durations specified in the figure legends. Sodium arsenite was used to induce oxidative stress, whereas cycloheximide (CHX; 10 μg/mL; Sigma-Aldrich, C4859-1ML) was used to inhibit translation elongation. The integrated stress response inhibitor ISRIB (250 nM; MedChemExpress, HY-12495A) was used to assess the contribution of eIF2α-dependent translational regulation to SG assembly. G3IB (50 μM; kindly provided by Paul Taylor’s laboratory)^21^ was used to inhibit SG formation. Actinomycin D (ActD; 5 μg/mL; supplier to be confirmed) was used to inhibit transcription in RNA decay experiments. Appropriate vehicle controls, including water, DMSO, or ethanol, were included as indicated.

For experiments involving pharmacological inhibition of SG assembly during hypoxia, G3IB was added 10 h after the initiation of hypoxic treatment, approximately 2 h before the appearance of SGs.

### Hypoxia treatment

U-2 OS cells were exposed to hypoxia (1% O₂) in a hypoxic chamber maintained at 37°C. Unless otherwise indicated, hypoxic treatments were performed for 12 h. All medium exchanges were performed using medium pre-equilibrated to 1% O₂. Cells were washed with pre-equilibrated PBS and subsequently fixed or lysed inside the hypoxic chamber to minimize reoxygenation during sample preparation.

### Surface sensing of translation (SUnSET) assay

Global protein synthesis was assessed using the surface sensing of translation (SUnSET) assay^22^. U-2 OS cells were seeded in 10 cm dishes and subjected to the indicated stress treatments. Puromycin (5 μg/mL; Thermo Fisher Scientific, A1113803) was added to the culture medium 15 min before the end of each treatment to label nascent polypeptides. Cells were subsequently harvested and lysed using RIPA buffer. Puromycin incorporation into newly synthesized proteins was assessed by immunoblotting using an anti-puromycin antibody. Relative protein synthesis was determined by quantifying puromycin incorporation and normalizing to the corresponding loading control.

### Stress granule isolation and transcriptome analysis

Stress granules were isolated from U-2 OS cells expressing G3BP1–eGFP using fluorescence-activated non-membrane condensate isolation (FANCI)^23^, as previously described, with modifications. Cells were seeded in 10 cm dishes and subjected to the indicated stress treatments. Following treatment, the culture medium was removed and replaced with 8 mL of fresh culture medium containing 1% formaldehyde (Sigma-Aldrich, F8775-500ML). Cells were fixed for 15 min, after which fixation was quenched with 200 mM glycine (Sigma-Aldrich, G7126-500G) for 5 min.

Cells were washed twice with ice-cold PBS and lysed in 2.1 mL of B0 buffer containing 50 mM HEPES (pH 7.5; Sigma-Aldrich, H0887-100ML), 150 mM KCl (Sigma-Aldrich, 60135-250ML), 1% NP-40 (Thermo Fisher Scientific, 85124), protease/phosphatase inhibitor cocktail (Cell Signaling Technology, 5872S), 1 mM dithiothreitol (DTT; Thermo Fisher Scientific, R0861), and RNase inhibitor (New England Biolabs, M0314L).

Cells were collected by scraping, and the lysates were divided equally among three 1.5 mL microcentrifuge tubes. Lysates were incubated on ice for 20 min before sonication using a Bioruptor Plus (Diagenode) for two cycles of 30 s ON and 30 s OFF at low power. Following sonication, lysates were incubated on ice for an additional 20 min, vortexed for 5 s, and passed through a 40 μm cell strainer to remove large cellular debris.

An aliquot of 50 μL was reserved as the whole-cell input for RNA-seq analysis, whereas the remaining lysate was subjected to fluorescence-activated sorting using a BD FACSAria III cell sorter (BD Biosciences) at the CSI Flow Cytometry Facility. G3BP1–eGFP-positive SG particles were collected in 100 μL of B0 buffer supplemented with additional RNase inhibitor (1:100; New England Biolabs, M0314L). Isolated SGs were washed twice with B0 buffer by centrifugation at 2,000 × g for 15 min using a swing-out rotor.

For RNA extraction, SG and input samples were adjusted to a final volume of 50 μL. Samples were digested with proteinase K (Sigma-Aldrich, 39450-01-6) for 40 min at 42°C with shaking at 500 rpm, followed by reversal of formaldehyde crosslinks at 65°C for 40 min without shaking. Samples were subsequently treated with TURBO DNase (Thermo Fisher Scientific, AM2238), and RNA was purified using the Oligo Clean & Concentrator Kit (Zymo Research, D4060). Purified RNA was submitted to Novogene for library preparation and RNA sequencing.

Three independent biological replicates were analyzed for each stress condition.

### Whole-cell mRNA sequencing

Whole-cell mRNA sequencing was performed to investigate SG-dependent changes in transcript abundance during hypoxia and RNA decay following transcriptional inhibition, as shown in Figures 6 and 7.

For hypoxic time-course experiments, U-2 OS cells expressing Halo–USP10p(1– 40)(F10A) or Halo–USP10p(1–40) were exposed to hypoxia (1% O₂) for 0, 6, 12, 18, or 24 h. For RNA decay experiments, U-2 OS cells were exposed to hypoxia for 12 h, with DMSO or G3IB added at 10 h. ActD (5 μg/mL) was added at 12 h, and cells were harvested at 0, 4, and 12 h following transcriptional inhibition.

Following treatment, cells were washed once with 1× PBS and lysed directly in 1× DNA/RNA Shield. Samples were submitted to Plasmidsaurus for Ultrafast RNA sequencing.

Three independent biological replicates were analysed per experimental condition.

### RNA sequencing and bioinformatic analysis

RNA sequencing was performed to characterize the transcriptomes of hypoxia- and arsenite-induced SGs and to investigate SG-dependent changes in RNA abundance and stability during hypoxia.

For SG transcriptome analysis, RNA isolated from purified SGs and their corresponding whole-cell inputs was subjected to library preparation and high-throughput sequencing by Novogene. For whole-cell transcriptome analysis, RNA samples collected from the indicated hypoxic time-course and Actinomycin D chase experiments were subjected to Ultrafast RNA sequencing by Plasmidsaurus.

Sequencing reads were processed and aligned to the human reference transcriptome or genome, followed by gene-level quantification. RNA abundance was normalized to account for differences in sequencing depth and library composition. Differential expression analyses were performed to identify transcripts exhibiting significant changes in abundance between experimental conditions.

For SG transcriptome analysis, RNA enrichment was calculated by comparing transcript abundance in isolated SGs with the corresponding whole-cell input and expressed as log₂ fold change (SG/input). Transcripts were classified as SG-enriched or SG-depleted based on their differential enrichment and posterior probability of differential expression (PPEE), using a threshold of 0.05.

To identify SG-dependent changes in RNA abundance during hypoxia, transcript abundance was compared between SG-competent U-2 OS cells expressing USP10p^1–40(F10A)^ and SG-deficient cells expressing USP10p^(1–40)^. For time-course analyses, RNA abundance was log₂-transformed, and changes at each time point were calculated relative to the corresponding 0 h condition independently in each cell line. The resulting log₂ fold changes were compared between SG-competent and SG-deficient cells to calculate the ΔΔ log₂ fold change. Transcripts exhibiting significant differential regulation at 12, 18, or 24 h of hypoxia (adjusted P < 0.05) were selected for downstream functional analyses.

For ActD chase experiments, RNA abundance was compared between DMSO- and G3IB-treated cells at the indicated time points following transcriptional inhibition. Transcript abundance was normalized to the corresponding 0 h ActD condition to assess relative RNA decay.

Gene Ontology (GO) enrichment analyses were performed on the indicated transcript sets to identify overrepresented biological processes, molecular functions, and cellular components. GO terms with a false discovery rate (FDR) below 0.05 were considered significantly enriched.

### Immunofluorescence microscopy

U-2 OS cells were seeded onto ethanol-treated No. 1.5 glass coverslips and subjected to the indicated stress treatments. Following treatment, cells were washed with 1× PBS and fixed with 4% paraformaldehyde (PFA) for 5 min. For hypoxic experiments, fixation was performed inside the hypoxic chamber to prevent reoxygenation.

Cells were permeabilized with 1% Triton X-100 in PBS for 5 min, followed by two washes with PBS. Cells were incubated with the indicated primary antibodies for 1 h at room temperature, washed three times with PBS, and subsequently incubated with the appropriate fluorescently labeled secondary antibodies for 1 h at room temperature. Following three additional washes with PBS, coverslips were mounted onto glass slides using mounting medium (Thermo Fisher Scientific, P36981).

Fluorescence images were acquired using a Zeiss Axio Imager M2 wide-field fluorescence microscope at the Microscopy and Multiplex Assays Core, CSI. Antibodies used for immunofluorescence are listed in Table 3.

**Table 3.**
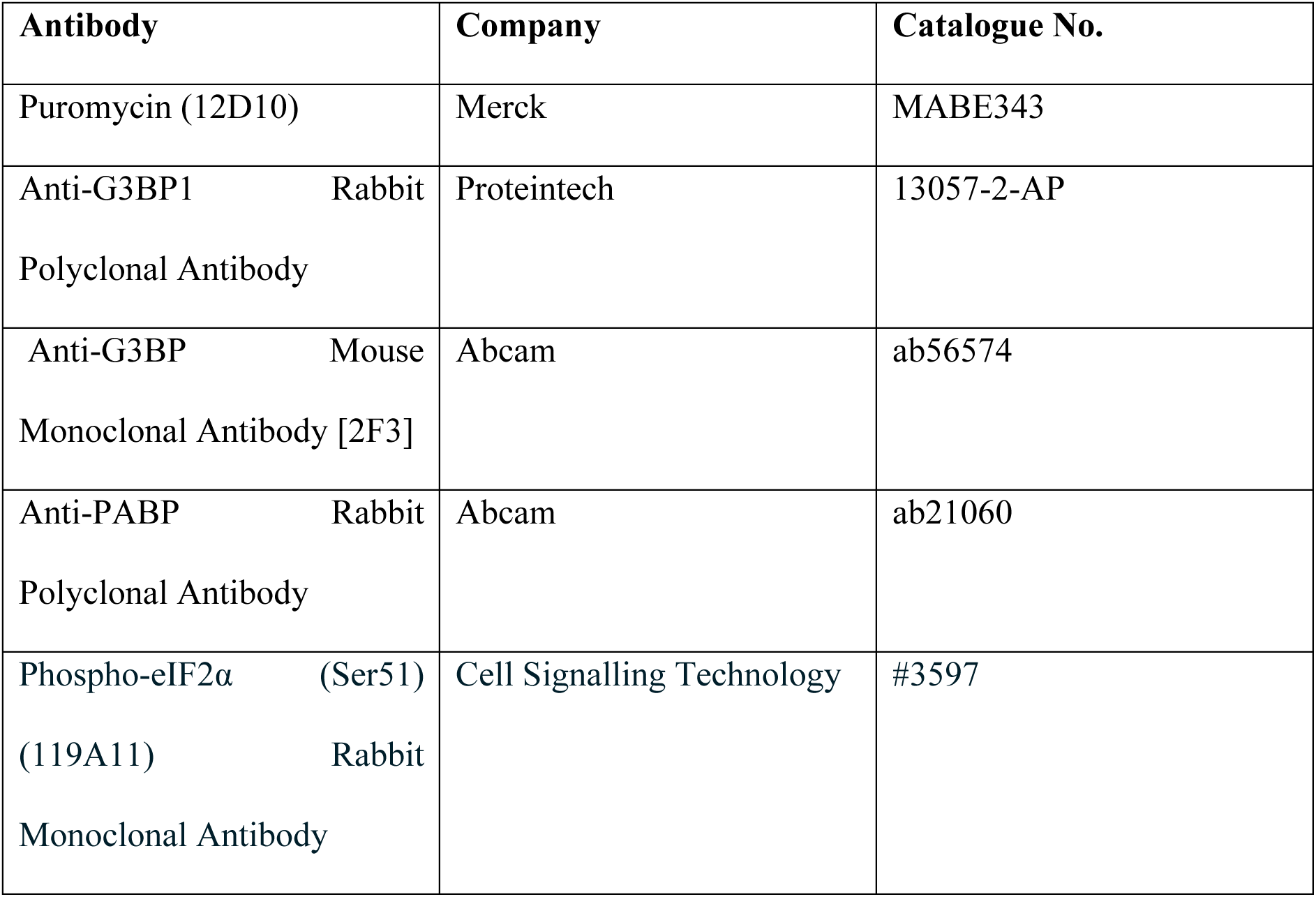
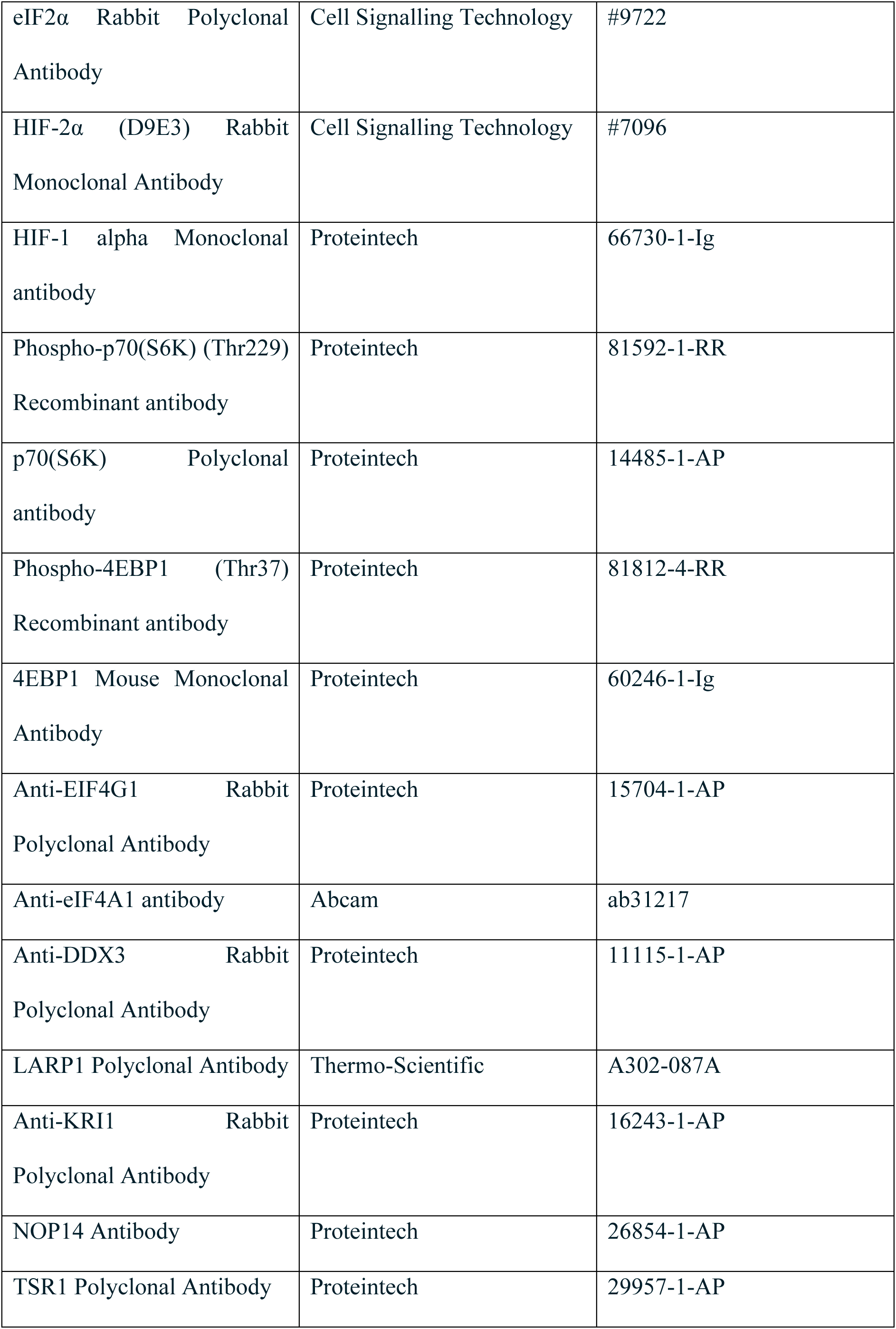
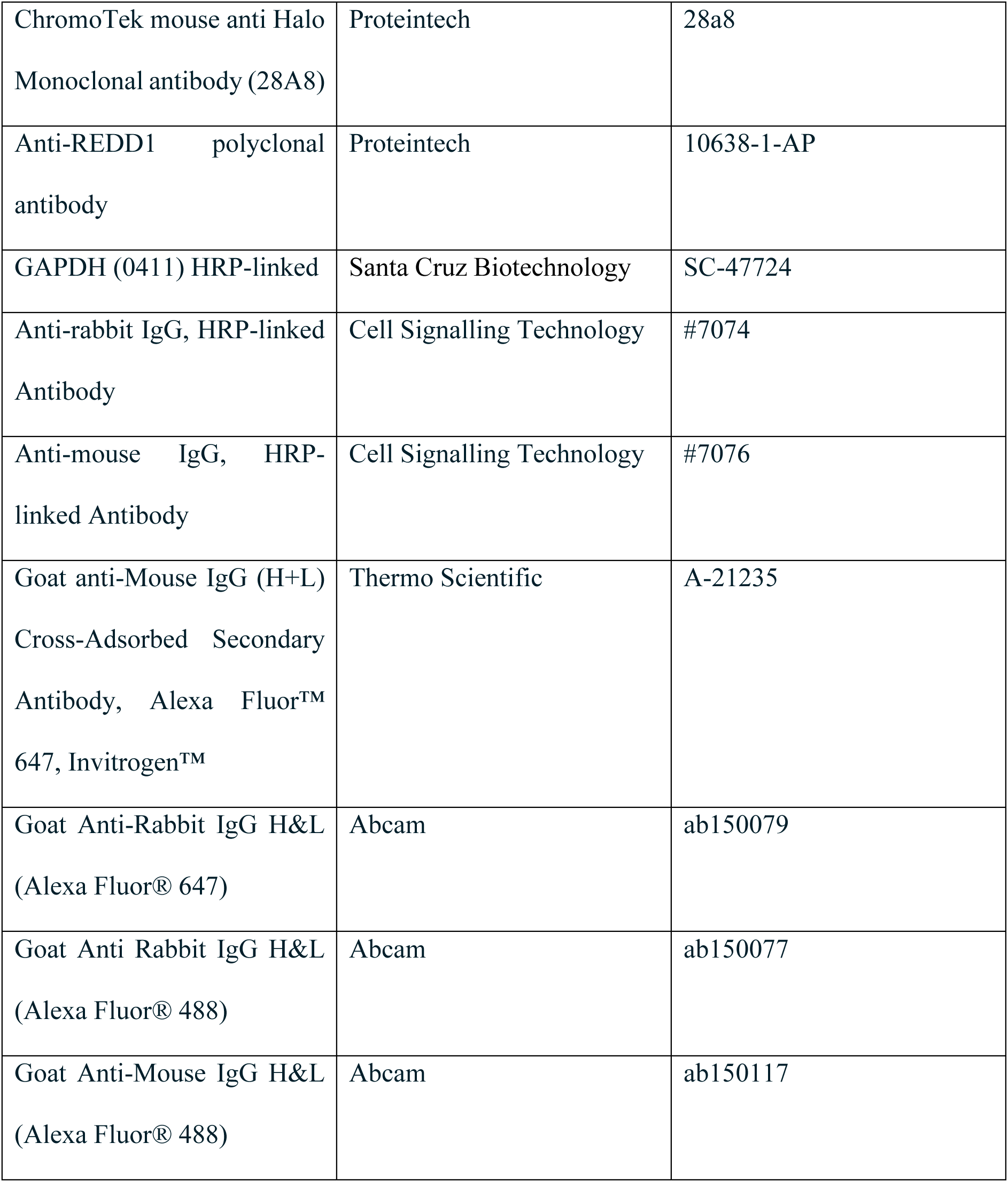
Antibodies used for western blotting and immunofluorescence.

| **Antibody** | **Company** | **Catalogue No.** |
|---|---|---|
| Puromycin (12D10) | Merck | MABE343 |
| Anti-G3BP1 Rabbit Polyclonal Antibody | Proteintech | 13057-2-AP |
| Anti-G3BP Mouse Monoclonal Antibody [2F3] | Abcam | ab56574 |
| Anti-PABP Rabbit Polyclonal Antibody | Abcam | ab21060 |
| Phospho-eIF2α (Ser51) (119A11) Rabbit Monoclonal Antibody | Cell Signalling Technology | #3597 |
| eIF2α Rabbit Polyclonal Antibody | Cell Signalling Technology | #9722 |
| HIF-2α (D9E3) Rabbit Monoclonal Antibody | Cell Signalling Technology | #7096 |
| HIF-1 alpha Monoclonal antibody | Proteintech | 66730-1-Ig |
| Phospho-p70(S6K) (Thr229) Recombinant antibody | Proteintech | 81592-1-RR |
| p70(S6K) Polyclonal antibody | Proteintech | 14485-1-AP |
| Phospho-4EBP1 (Thr37) Recombinant antibody | Proteintech | 81812-4-RR |
| 4EBP1 Mouse Monoclonal Antibody | Proteintech | 60246-1-Ig |
| Anti-EIF4G1 Rabbit Polyclonal Antibody | Proteintech | 15704-1-AP |
| Anti-eIF4A1 antibody | Abcam | ab31217 |
| Anti-DDX3 Rabbit Polyclonal Antibody | Proteintech | 11115-1-AP |
| LARP1 Polyclonal Antibody | Thermo-Scientific | A302-087A |
| Anti-KRI1 Rabbit Polyclonal Antibody | Proteintech | 16243-1-AP |
| NOP14 Antibody | Proteintech | 26854-1-AP |
| TSR1 Polyclonal Antibody | Proteintech | 29957-1-AP |
| ChromoTek mouse anti Halo Monoclonal antibody (28A8) | Proteintech | 28a8 |
| Anti-REDD1 polyclonal antibody | Proteintech | 10638-1-AP |
| GAPDH (0411) HRP-linked | Santa Cruz Biotechnology | SC-47724 |
| Anti-rabbit IgG, HRP-linked Antibody | Cell Signalling Technology | #7074 |
| Anti-mouse IgG, HRP-linked Antibody | Cell Signalling Technology | #7076 |
| Goat anti-Mouse IgG (H+L) Cross-Adsorbed Secondary Antibody, Alexa Fluor™ 647, Invitrogen™ | Thermo Scientific | A-21235 |
| Goat Anti-Rabbit IgG H&L (Alexa Fluor® 647) | Abcam | ab150079 |
| Goat Anti Rabbit IgG H&L (Alexa Fluor® 488) | Abcam | ab150077 |
| Goat Anti-Mouse IgG H&L (Alexa Fluor® 488) | Abcam | ab150117 |

### Immunoprecipitation

GFP-Trap immunoprecipitation was performed to assess the association of G3BP1 with LARP1 and other SG-associated proteins. U-2 OS cells expressing G3BP1–GFP were subjected to the indicated stress treatments and lysed in immunoprecipitation buffer containing NaCl, Tris-HCl (pH 7.5), EDTA, NP-40, glycerol, and protease/phosphatase inhibitors (Cell Signaling Technology, 5872S). GFP-tagged protein complexes were isolated using GFP-Trap magnetic agarose beads (ChromoTek, gtma-20) according to the manufacturer’s instructions.

For experiments assessing RNA-dependent protein associations, cell lysates were treated with RNase A before immunoprecipitation. Immunoprecipitated complexes were subsequently analyzed by immunoblotting using antibodies against G3BP1, LARP1, and CAPRIN1, as indicated.

### Single-molecule fluorescence in situ hybridization (smFISH) and image analysis

Single-molecule fluorescence in situ hybridization (smFISH) was performed using a protocol adapted from Khong et al. (2018), with modifications^24,25^. Briefly, fixed U-2 OS cells were permeabilized with 1% Triton X-100 in PBS for 5 min. Cells were subsequently processed for hybridization with fluorescently labeled oligonucleotide probes targeting the indicated transcripts.

Following hybridization and washing, cells were immunostained for G3BP1 to identify SGs and counterstained with DAPI to visualize nuclei. Fluorescence images were acquired using Live-SR spinning disc confocal with TIRF (CBIS, NUS) and analyzed using Imaris image analysis software (Bitplane/Oxford Instruments at CBIS, NUS).

Individual RNA molecules were identified using the Imaris Spots tool, whereas G3BP1-positive SGs were identified using fluorescence-based segmentation (Imaris Surface Tool). To determine the fraction of individual transcripts localized within SGs, the number of smFISH spots overlapping G3BP1-positive SGs was divided by the total number of smFISH spots detected within each cell.

The fraction of RNA localized within SGs was calculated as: Fraction of RNA in SGs = Number of RNA spots overlapping SGs / Total cytosolic number of RNA spots per cell.

### Western blotting

Cells were washed once with ice-cold PBS and lysed using RIPA buffer (Thermo Fisher Scientific, 89990) supplemented with 1× protease/phosphatase inhibitor cocktail (Cell Signaling Technology, 5872S). Cells were scraped from culture dishes, and lysates were incubated on ice for 10 min. Insoluble cellular debris was removed by centrifugation, and the clarified lysates were collected for protein quantification.

Protein lysates were mixed with 4× Bolt LDS Sample Buffer (Thermo Fisher Scientific, B0007) and 10× Bolt Sample Reducing Agent (Thermo Fisher Scientific, B0009). Samples were denatured at 95°C for 5 min before separation by SDS–PAGE using Bolt gels (Thermo Fisher Scientific, NW04120BOX). Electrophoresis was performed at 200 V for 20 min in 1× Bolt MOPS SDS Running Buffer (Thermo Fisher Scientific, B000102).

Proteins were transferred onto nitrocellulose membranes using the iBlot 3 transfer system (Thermo Fisher Scientific, IB31001). Membranes were blocked for 30 min in either 5% bovine serum albumin (BSA; Sigma-Aldrich, A6003) in Tris-buffered saline containing Tween-20 (TBST) for phosphoprotein detection or 5% non-fat milk in TBST for other proteins. Membranes were incubated with the indicated primary antibodies diluted in the corresponding blocking buffer for 1 h at room temperature with continuous agitation. Following three washes with TBST, membranes were incubated with the appropriate secondary antibodies for 1 h at room temperature. Membranes were washed three additional times with TBST, and immunoreactive proteins were detected using Pierce ECL Western Blotting Substrate (Thermo Fisher Scientific, 32106).

Where required, membranes were stripped using Restore PLUS Western Blot Stripping Buffer (Thermo Fisher Scientific, 46430) for 15 min at room temperature before reblocking and subsequent antibody probing.

Antibodies used for western blotting listed in Table 3.

### Cell viability and metabolic activity assay (CCK-8)

Cellular metabolic activity was assessed using the Cell Counting Kit-8 (CCK-8) assay. U-2 OS cells were seeded in 96-well plates and allowed to adhere overnight before exposure to hypoxia (1% O₂) for the indicated durations. To assess the contribution of SGs to cellular metabolic activity during hypoxia, cells were treated with either G3IB (50 μM) or an equivalent volume of DMSO as a vehicle control. G3IB was added 10 h after the initiation of hypoxia, approximately 2 h before SG assembly.

At the indicated time points, CCK-8 reagent was added to each well according to the manufacturer’s instructions, and cells were incubated to allow development of the colorimetric signal. Absorbance was measured at 450 nm using a microplate reader. Background absorbance from medium-only wells was subtracted, and values were normalized to the corresponding 0 h normoxic DMSO-treated controls.

Experiments were performed using three independent biological replicates (n = 3), and data are presented as mean ± SD. Statistical significance was determined using unpaired Student’s t-tests.

### Cell cycle analysis by flow cytometry

Cell cycle distribution was assessed by flow cytometry using DAPI staining to quantify cellular DNA content. U-2 OS cells were subjected to the indicated stress treatments, harvested by trypsinization, and collected by centrifugation at 300 × g for 5 min. Cells were washed and resuspended in 1 mL of ice-cold PBS, followed by the gradual addition of 9 mL of ice-cold methanol for fixation. Fixed cells were incubated at 4°C for at least 1 h and subsequently collected by centrifugation at 300 × g for 5 min. Cell pellets were resuspended in PBS containing DAPI (1:1,000) and incubated for 15 min at 4°C with agitation.

Cellular DNA content was measured using a BD LSRFortessa X-20 flow cytometer (BD Biosciences) at the Flow Cytometry Facility, Cancer Science Institute of Singapore (CSI). Cell debris and doublets were excluded during flow cytometric analysis, and cell cycle distributions were determined based on DAPI fluorescence intensity. Cells were classified into G0/G1, S, and G2/M phases according to DNA content.

Experiments were performed using three independent biological replicates (n = 3), and data are presented as mean ± SD. Statistical significance was determined using unpaired Student’s t-tests.

### Statistical analysis

Statistical analyses were performed using the indicated tests, with three independent biological replicates (n = 3) unless otherwise stated. Data are presented as mean ± standard deviation (SD).

For comparisons between two experimental conditions, statistical significance was determined using two-sided unpaired Student’s t-tests. For RNA abundance time-course experiments, mixed-effects models fitted by restricted maximum likelihood (REML) were used to assess the interaction between hypoxia duration and SG competence (time × cell line). Comparisons between SG-competent and SG-deficient cells at individual time points were performed using unpaired Student’s t-tests.

For SG transcriptome analyses, differential RNA enrichment was assessed using posterior probabilities of differential expression (PPEE), with a threshold of 0.05 used to identify significantly enriched or depleted transcripts.

For whole-cell RNA-seq analyses, SG-dependent differential regulation was evaluated by comparing changes in log₂-transformed RNA abundance relative to baseline (0 h) between SG-competent and SG-deficient cells (ΔΔ log₂ fold change). Uncertainty associated with ΔΔ log₂ fold changes was estimated by propagating the standard deviations of the underlying log₂-transformed measurements. Specifically, the variances associated with the 0 h and corresponding hypoxic time point were summed for each cell line, followed by summation of the variances from SG-competent and SG-deficient cells. The propagated standard deviation was calculated as the square root of the total variance, assuming independent measurements.

Transcripts with adjusted P values below 0.05 were considered significantly differentially regulated. The statistical test used to calculate P values and the multiple-testing correction method remain to be confirmed.

Statistical significance is indicated as follows: ns, not significant; *P < 0.05, **P < 0.01, ***P < 0.001, and ****P < 0.0001. Exact statistical tests and significance thresholds are indicated in the corresponding figure legends.

## RESULTS

### Hypoxia induces stress granule formation without substantially compromising cell survival

To investigate stress granule (SG) biology under hypoxic conditions, we exposed U-2 OS cells to 1% O₂, an oxygen concentration commonly encountered within tumour microenvironments^26^, for varying durations. Immunofluorescence analysis of the canonical SG marker G3BP1 revealed robust SG formation following 12h of hypoxic exposure (Figure 1A–B).

**Fig 1.**
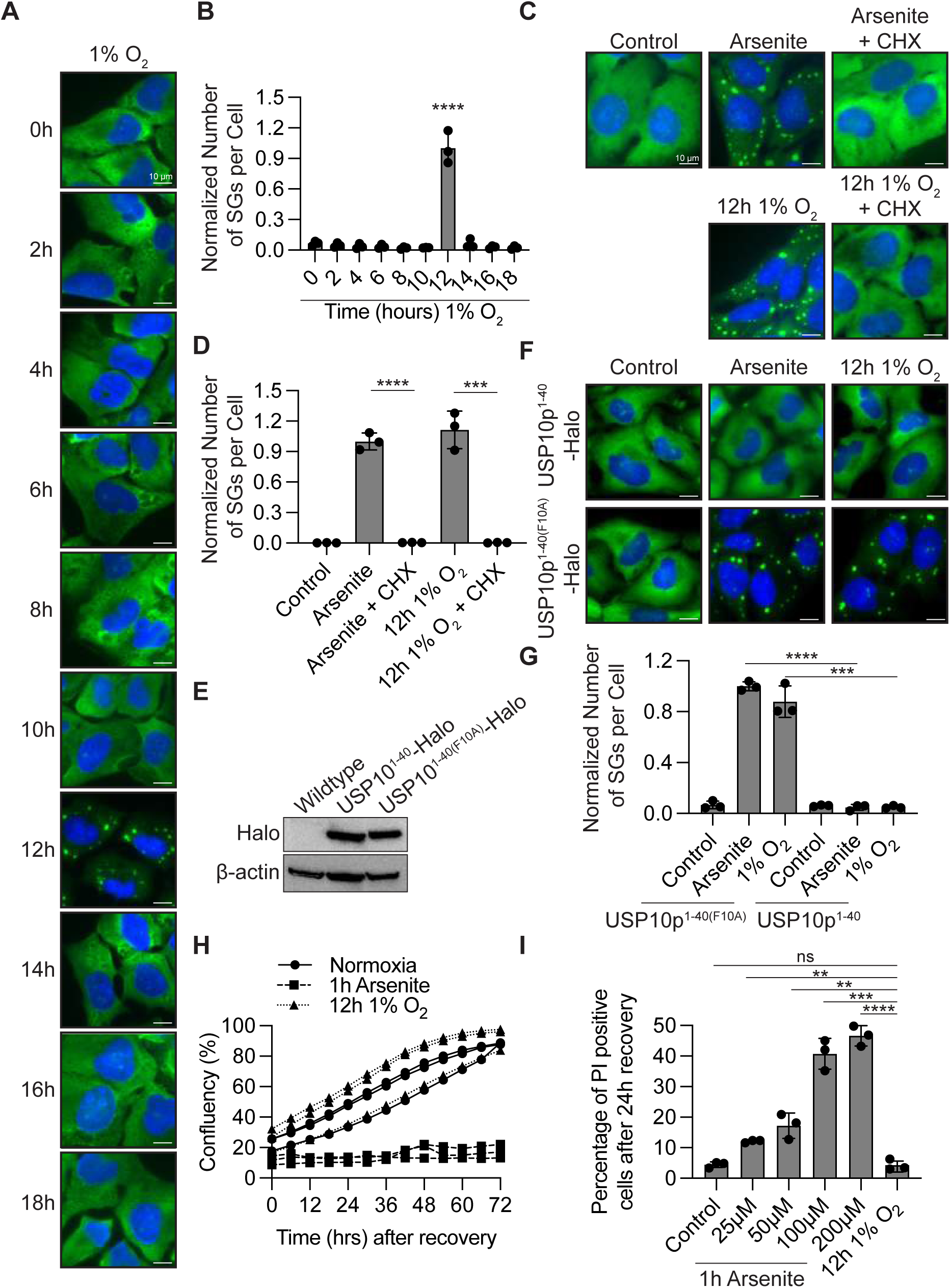
Hypoxia induces transient stress granule assembly with minimal cytotoxicity. **(A)** Representative immunofluorescence images of U-2 OS cells exposed to hypoxia (1% O_2_) for the indicated durations. SGs were visualized by G3BP1 immunostaining (green), and nuclei were counterstained with DAPI (blue). Scale bars, 10 µm. **(B)** Quantification of SGs per cell from (A), normalized to arsenite-treated controls. **(C)** Representative immunofluorescence images of U-2 OS cells treated with arsenite or exposed to hypoxia (1% O_2_, 12 h), in the presence or absence of (CHX; duration last two hours and last hour respectively). SGs were visualized by G3BP1 immunostaining (green), and nuclei were counterstained with DAPI (blue). Scale bars, 10 µm. **(D)** Quantification of SGs per cell from (A), normalized to arsenite-treated controls. **(E)** Immunoblot analysis of Halo-tagged USP10p^1–40^ and its F10A mutant, USP10p^1–40(F10A)^, expressed in U-2 OS cells. β-actin serves as a loading control. **(F)** Representative immunofluorescence images of U-2 OS cells expressing USP10p^1–40^ or USP10p^1–40(F10A)^ under unstressed conditions, arsenite treatment, or hypoxia (1% O_2_, 12 h). SGs were visualized by G3BP1 immunostaining (green), and nuclei were counterstained with DAPI (blue). Scale bars, 10 µm. **(G)** Quantification of SGs per cell from (F), normalized to arsenite-treated controls. **(H)** Cell proliferation measured by live-cell confluency over 72h following recovery from arsenite treatment (1h) or hypoxia (1% O_2_, 12h). Following stress exposure, arsenite-treated cells were transferred to arsenite-free medium, whereas hypoxia-treated cells were returned to normoxic conditions. **(I)** Quantification of percentages of propidium iodide (PI)-positive U-2 OS cells following 24 h o f recovery from arsenite treatment (25-200 µM, 1 h) or hypoxia (1% O_2_, 12 h). Data are presented as mean ± SD from three independent biological replicates (n = 3). Statistical significance was determined using unpaired Student’s t-tests. Ns, not significant; ** P < 0;01, *** P < 0.001, **** P < 0.0001.

Because hypoxia-induced SGs remain less extensively characterized than those formed in response to classical stressors, we first asked whether their assembly follows the established principles of canonical SG formation. A defining feature of SG assembly is the accumulation of non-translating messenger ribonucleoprotein complexes (mRNPs), which provide the molecular substrates for condensation^7,27^. To determine whether hypoxia-induced SGs similarly depend on non-translating mRNPs, we treated cells with cycloheximide after 10 h of hypoxic exposure, prior to detectable SG formation, and assessed SG assembly 2 h later. Cycloheximide treatment effectively prevented hypoxia-induced SG formation (Figure 1C–D), indicating that the availability of non-translating mRNPs is required for SG assembly under hypoxic conditions.

We next examined whether hypoxia-induced SG formation depends on G3BP-mediated multivalent interactions^10–12^, another defining feature of canonical SG assembly. To disrupt these interactions, we stably expressed a peptide corresponding to the N-terminal 40 amino acids of USP10 (USP10p^1–40^), which binds G3BP1/2 and inhibits SG assembly by interfering with G3BP-dependent protein interactions and multivalency^28,29^. Expression of USP10p^1–40^ strongly suppressed SG formation under hypoxia, whereas expression of the binding-deficient USP10p^1–40(F10A)^ mutant did not (Figure 1E–G). Together, the sensitivity of hypoxia-induced granules to cycloheximide and disruption of G3BP-mediated interactions supports their classification as bona fide SGs.

Having established that hypoxia induces SG assembly, we next asked whether SG formation occurs under conditions that substantially compromise cellular viability. U-2 OS cells were exposed to 1% O_2_ for 12 h, followed by reoxygenation under normoxic conditions. Cellular recovery was assessed by monitoring confluency using CellCyte, while cell death was evaluated by propidium iodide (PI) staining. Cells previously exposed to hypoxia resumed proliferation at rates comparable to those maintained continuously under normoxia (Figure 1H). Consistent with this observation, hypoxia-exposed cells exhibited minimal PI staining (Figure 1I). In contrast, cells exposed to SG-inducing concentrations of sodium arsenite displayed little to no proliferative recovery following stress removal and substantially increased PI staining (Figure 1H–I).

Collectively, these findings demonstrate that prolonged exposure to 1% O_2_ induces bona fide SGs without substantially compromising cell survival, in contrast to the pronounced cytotoxicity associated with arsenite treatment. Thus, hypoxia provides a comparatively non-cytotoxic physiological stress condition in which to investigate the mechanisms and functions of SG assembly.

### Hypoxia-induced stress granules assemble through an ISR-independent HIF1α–REDD1– mTORC1–4E-BP1 signalling axis

Having established that hypoxia induces bona fide SGs, we next sought to identify the signalling mechanisms responsible for their assembly. Under many stress conditions, SG formation is driven by activation of the integrated stress response (ISR), in which phosphorylation of eIF2α inhibits translation initiation and promotes the accumulation of non-translating mRNPs^8,30,31^. We therefore asked whether hypoxia-induced SG assembly similarly depends on ISR activation.

Immunoblot analysis revealed little to no detectable increase in eIF2α phosphorylation following exposure to 1% O_2_, in marked contrast to arsenite treatment, which robustly induced eIF2α phosphorylation under SG-forming conditions (Figure 2A). These findings suggested that hypoxia-induced SG formation may occur independently of substantial ISR activation. However, we considered the possibility that low levels of eIF2α phosphorylation, below the detection threshold of immunoblotting, could nevertheless contribute to SG assembly.

**Fig 2.**
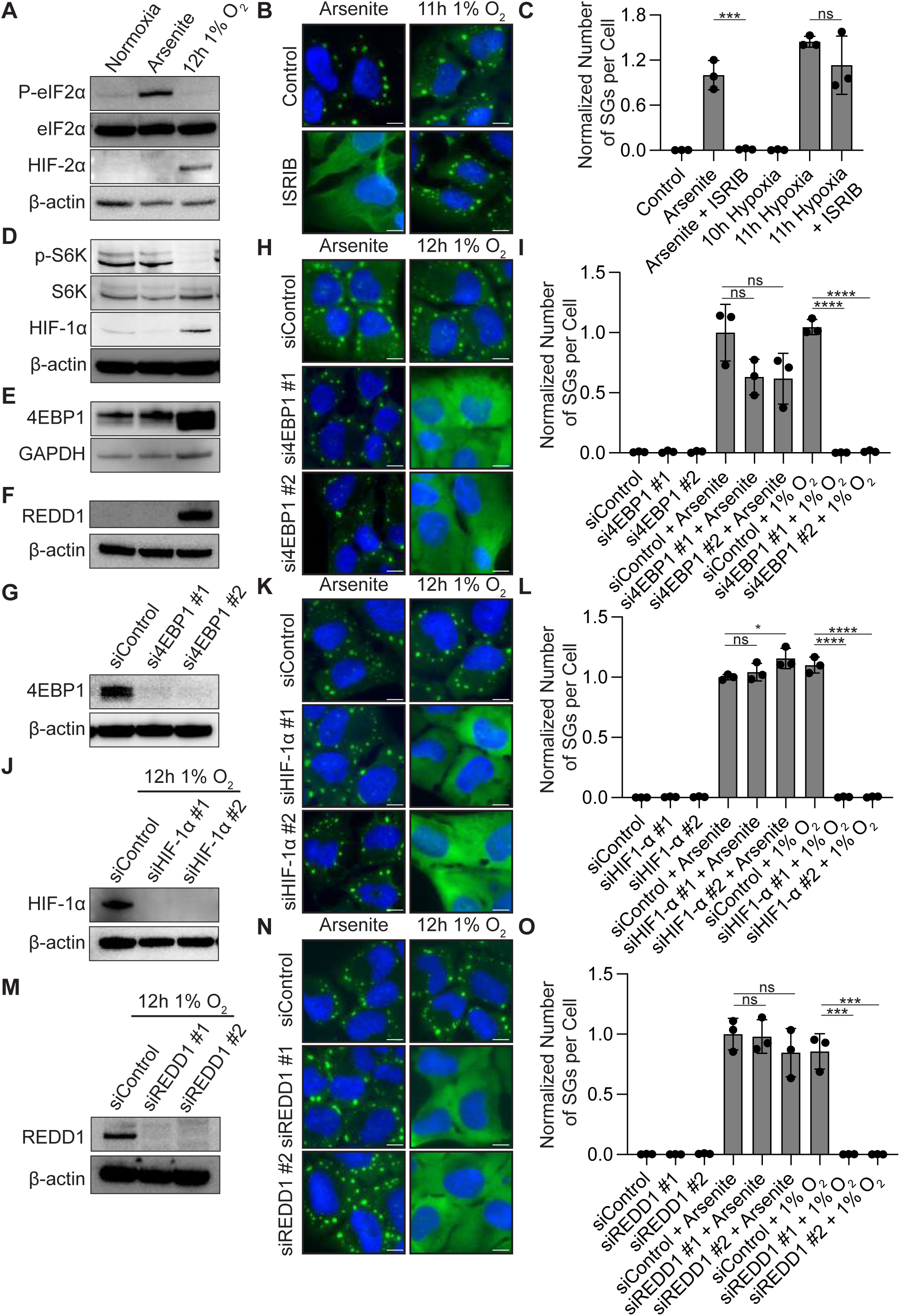
Hypoxia-induced stress granule assembly is independent of the integrated stress response but requires 4E-BP1. **(A)** Immunoblot analysis of phosphorylated eIF2α (P-eIF2α), total eIF2α, HIF-2α, and β-actin in U-2 OS cells maintained under normoxia, treated with arsenite, or exposed to hypoxia (1% O_2_, 12 h). **(B)** Representative immunofluorescence images of U-2 OS cells treated with 1 h arsenite or exposed to hypoxia (1% O_2_, 11 h), in the presence or absence of the integrated stress response inhibitor ISRIB for the last hour. SGs were visualized by G3BP1 immunostaining (green), and nuclei were counterstained with DAPI (blue). Scale bars, 10 μm. **(C)** Quantification of SGs per cell under the indicated conditions in (B), normalized to arsenite-treated controls. **(D–F)** Immunoblot analysis of phosphorylated S6K (P-S6K), total S6K, HIF-1α, 4E-BP1, and REDD1 in U-2 OS cells maintained under normoxia, treated with arsenite, or exposed to hypoxia (1% O_2_, 12 h). β-actin or GAPDH serves as the loading control. **(G)** Immunoblot analysis confirming siRNA-mediated depletion of 4E-BP1 using two independent siRNAs (si4EBP1 #1 and #2) compared with a non-targeting control (siControl). β-actin serves as a loading control. **(H)** Representative immunofluorescence images of U-2 OS cells transfected with siControl or two independent siRNAs targeting 4E-BP1, followed by 1 h arsenite treatment or hypoxia (1% O_2_, 12 h). SGs were visualized by G3BP1 immunostaining (green), and nuclei were counterstained with DAPI (blue). Scale bars, 10 μm. **(I)** Quantification of SGs per cell from (H), normalized to arsenite-treated controls. **(J)** Immunoblot analysis confirming siRNA-mediated depletion of HIF-1α using two independent siRNAs (siHIF-1α #1 and #2) following hypoxia (1% O_2_, 12 h). β-actin serves as a loading control. **(K)** Representative immunofluorescence images of U-2 OS cells transfected with siControl or two independent siRNAs targeting HIF-1α, followed by arsenite treatment or hypoxia (1% O_2_, 12 h). SGs were visualized by G3BP1 immunostaining (green), and nuclei were counterstained with DAPI (blue). Scale bars, 10 μm. **(L)** Quantification of SGs per cell from (K), normalized to arsenite-treated controls. **(M)** Immunoblot analysis confirming siRNA-mediated depletion of REDD1 using two independent siRNAs (siREDD1 #1 and #2) following hypoxia (1% O_2_, 12 h). β-actin serves as a loading control. **(N)** Representative immunofluorescence images of U-2 OS cells transfected with siControl or two independent siRNAs targeting REDD1, followed by arsenite treatment or hypoxia (1% O_2_, 12 h). SGs were visualized by G3BP1 immunostaining (green), and nuclei were counterstained with DAPI (blue). Scale bars, 10 μm. **(O)** Quantification of SGs per cell from (N), normalized to arsenite-treated controls. Data are presented as mean ± SD from three independent biological replicates (n = 3). Statistical significance was determined using unpaired Student’s t-tests. ns, not significant; *P < 0.05, ***P < 0.001, ****P < 0.0001.

To functionally assess ISR dependence, we treated cells with ISRIB, a small-molecule activator of eIF2B that counteracts ISR-mediated translational repression^32–34^. Consistent with previous report^35^, ISRIB effectively suppressed arsenite-induced SG formation (Figure 2B–C). In contrast, ISRIB had little effect on hypoxia-induced SG assembly (Figure 2B–C). Together, these observations indicate that hypoxia-induced SG formation does not require canonical ISR-mediated translational repression, suggesting that an alternative mechanism promotes the accumulation and condensation of non-translating mRNPs.

We next considered whether hypoxia-induced SG assembly might instead involve inhibition of mechanistic target of rapamycin complex 1 (mTORC1), another major regulator of translation initiation. Hypoxia is known to suppress mTORC1 activity, in part through HIF-dependent induction of REDD1^36^. To examine this pathway, we assessed the phosphorylation of S6 kinase (S6K) and 4E-BP1, two established downstream readouts of mTORC1 activity, together with the expression of HIF1α and REDD1^37^. Exposure to 1% O_2_ resulted in a marked reduction in S6K phosphorylation and increased electrophoretic mobility of 4E-BP1, consistent with its hypophosphorylation (Figure 2D–E). Concurrently, hypoxia induced HIF1α and REDD1 expression (Figure 2A,F). These findings indicate that hypoxia suppresses mTORC1 signalling under conditions that promote SG assembly.

To determine whether this pathway contributes functionally to SG formation, we first examined the requirement for 4E-BP1, a translational repressor activated upon mTORC1 inhibition^38^. Hypophosphorylated 4E-BP1 binds eIF4E and prevents its association with eIF4G, thereby restricting eIF4F complex assembly and cap-dependent translation initiation^39,40^. Depletion of 4E-BP1 using two independent siRNAs strongly impaired hypoxia-induced SG formation while having minimal effect on arsenite-induced SG assembly (Figure 2G–I). These findings identify 4E-BP1 as a critical regulator of SG formation specifically under hypoxic conditions.

We next asked whether upstream components of the hypoxia–mTORC1 signalling pathway were similarly required for SG assembly. Under hypoxia, HIF1α transcription factor promote REDD1 expression^36^, which contributes to mTORC1 inhibition. Depletion of HIF1α markedly suppressed hypoxia-induced SG formation without detectably affecting arsenite-induced SGs (Figure 2J–L). Likewise, depletion of REDD1 strongly impaired SG assembly under hypoxia but not arsenite stress (Figure 2M–O).

Together, these findings establish that hypoxia-induced SG assembly depends on a HIF–REDD1–mTORC1–4E-BP1 signalling axis rather than canonical ISR activation. Thus, although hypoxia- and arsenite-induced SGs share common requirements for non-translating mRNPs and G3BP-mediated interactions, they arise through distinct upstream signalling mechanisms.

### Hypoxia-induced stress granules assemble despite sustained global translation

SG assembly is generally associated with extensive translational repression, which increases the availability of non-translating mRNPs capable of undergoing condensation. Given that hypoxia-induced SG assembly depends on a signalling pathway distinct from the canonical ISR, we next asked whether hypoxia similarly requires extensive translational repression to promote SG formation.

To address this, we measured global protein synthesis using the SUnSET^22^ assay under hypoxic and arsenite stress conditions. Surprisingly, global translational activity remained largely sustained during hypoxia-induced SG formation, in marked contrast to the pronounced translational repression observed following arsenite treatment (Figure 3A–B). These findings indicate that hypoxia-induced SGs can assemble without extensive global translational shutdown, raising the possibility that a comparatively limited pool of non-translating mRNPs is sufficient to support condensation under hypoxic conditions.

**Figure 3.**
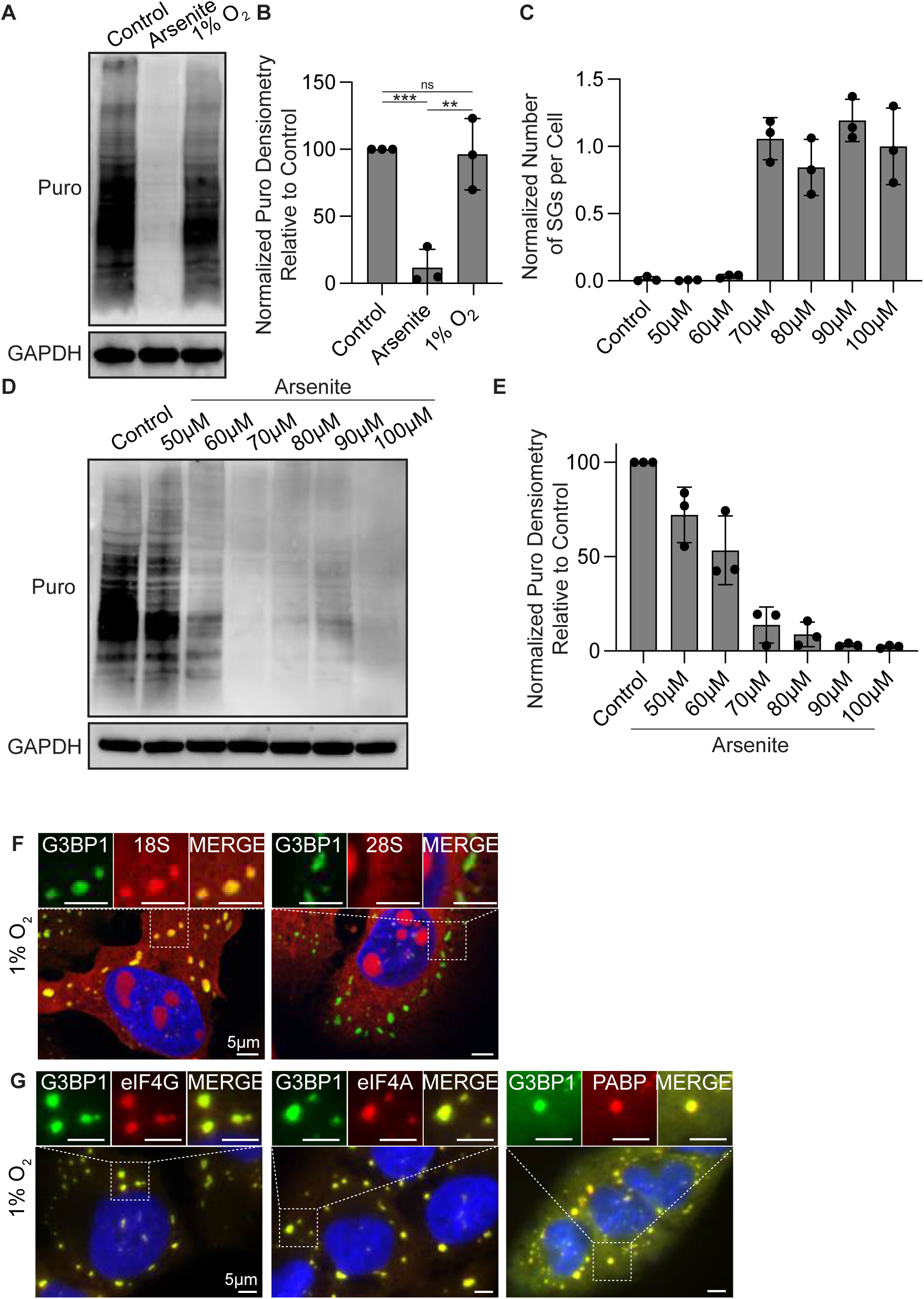
Hypoxia induces stress granule assembly with minimal global translational repression. **(A)** Representative immunoblot analysis of puromycin incorporation in U-2 OS cells maintained under normoxia (control), treated with arsenite (1 h), or exposed to hypoxia (1% O_2_, 12 h). Puromycin incorporation was measured using the SUnSET assay to assess global protein synthesis. GAPDH serves as a loading control. **(B)** Quantification of puromycin incorporation from (A), normalized to untreated controls. **(C)** Quantification of SGs per cell in U-2 OS cells treated with increasing concentrations of arsenite (50–100 μM, 1 h), normalized to arsenite-treated controls. **(D)** Representative immunoblot analysis of puromycin incorporation in U-2 OS cells treated with increasing concentrations of arsenite (50–100 μM, 1 h). GAPDH serves as a loading control. **(E)** Quantification of puromycin incorporation from (D), normalized to untreated controls. **(F)** Representative immunofluorescence and RNA fluorescence *in situ* hybridization (RNA-FISH) images of U-2 OS cells exposed to hypoxia (1% O_2_, 12 h), showing the localization of 18S or 28S ribosomal RNA (red) relative to G3BP1-positive SGs (green). Nuclei were counterstained with DAPI (blue). Enlarged images of the indicated regions are shown above. Scale bars, 5 μm. **(G)** Representative immunofluorescence images of U-2 OS cells exposed to hypoxia (1% O_2_, 12 h), showing the localization of translation initiation factors eIF4G and eIF4A, or poly(A)-binding protein (PABP) (red), relative to G3BP1-positive SGs (green). Nuclei were counterstained with DAPI (blue). Enlarged images of the indicated regions are shown above. Scale bars, 5 μm. Data are presented as mean ± SD from three independent biological replicates (n = 3). Statistical significance was determined using unpaired Student’s t-tests. ns, not significant; **P < 0.01, ***P < 0.001.

To determine whether arsenite-induced SGs could similarly assemble under conditions of modest translational repression, we titrated arsenite concentrations to identify the threshold required for SG formation. SGs were readily detected following treatment with 70 µM arsenite but were largely absent at 60 µM (Figure 3C). Notably, both concentrations caused substantial translational repression, as measured by SUnSET (Figure 3D–E). Thus, even arsenite concentrations approaching the threshold for SG assembly were associated with pronounced inhibition of protein synthesis. These observations are consistent with recent findings linking ISR-induced SG formation to substantial translational repression^41^.

Together, these results reveal a striking difference between hypoxia- and arsenite-induced SG assembly. Whereas arsenite-induced SG formation is associated with extensive global translational repression, hypoxia-induced SGs assemble despite relatively sustained protein synthesis. This suggests that hypoxia may engage additional mechanisms that facilitate condensation under conditions of comparatively modest translational inhibition.

Because global protein synthesis remained largely intact during hypoxia-induced SG assembly, we next considered whether actively translating ribosomes might contribute to the composition of these granules. Although cycloheximide sensitivity indicated that non-translating mRNPs are required for SG formation (Figure 1C–D), it remained possible that translating mRNPs or large ribosomal subunits could also partition into hypoxia-induced SGs. To investigate this possibility, we examined the localization of 18S and 28S rRNA, components of the small and large ribosomal subunits, respectively, using fluorescence in situ hybridization (Figure 3F). If actively translating ribosomes were substantially enriched within hypoxia-induced SGs, we would expect both ribosomal subunits to partition into these condensates. However, 28S rRNA was not detectably enriched within hypoxia-induced SGs, whereas 18S rRNA exhibited clear enrichment (Figure 3F). These observations are consistent with the accumulation of small ribosomal subunit-associated mRNPs rather than intact translating ribosomes.

We next asked whether hypoxia-induced SGs retain canonical translation initiation factors commonly associated with arsenite-induced SGs^8,42^. Immunofluorescence analysis revealed that eIF4G, eIF4A, and PABP were enriched within hypoxia-induced SGs at levels comparable to those observed during arsenite stress (Figure 3G). Thus, despite assembling under conditions of sustained global protein synthesis, hypoxia-induced SGs retain several characteristic components of canonical SGs.

Collectively, these findings demonstrate that hypoxia promotes the assembly of SGs containing small ribosomal subunit-associated mRNPs and canonical translation initiation factors without requiring extensive global translational repression. This raises the possibility that hypoxia engages additional molecular factors that facilitate SG condensation under conditions in which the availability of non-translating mRNPs may be comparatively limited.

### LARP1 selectively promotes stress granule assembly under hypoxic conditions

The ability of hypoxia-induced SGs to assemble despite relatively sustained global translation suggested that additional molecular factors might facilitate condensation under these conditions. Because 4E-BP1 was required for hypoxia-induced SG assembly, we first asked whether 4E-BP1 itself localizes to SGs and potentially contributes directly to their condensation. Surprisingly, immunofluorescence analysis revealed that 4E-BP1 was not detectably enriched within hypoxia-induced SGs (Figure 4A–B). These observations suggest that 4E-BP1 primarily promotes SG assembly through its regulation of translation initiation rather than by functioning as a major structural component of the condensates.

**Figure 4.**
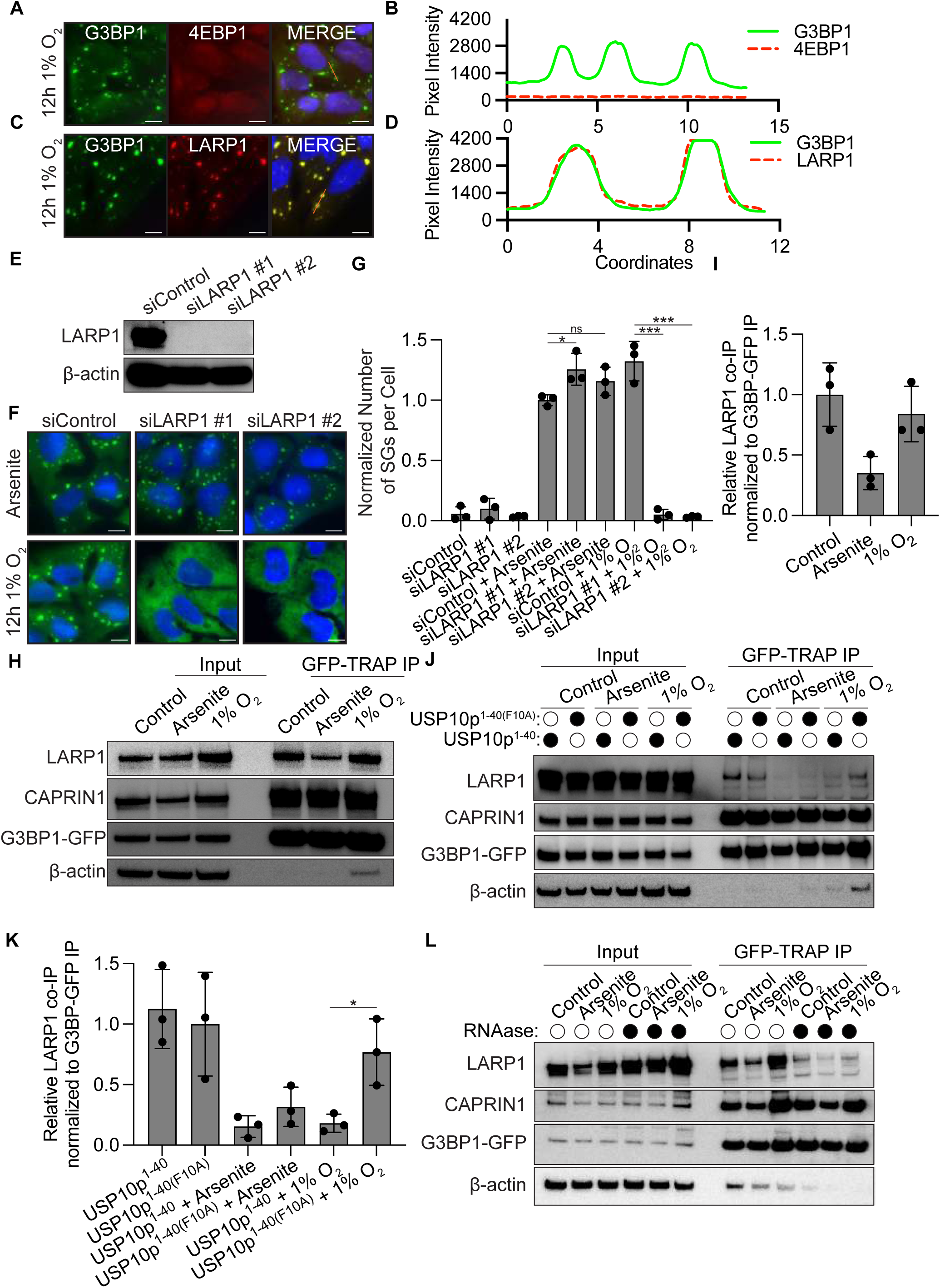
LARP1 associates with G3BP1 and is required for hypoxia-induced stress granule assembly. **(A)** Representative immunofluorescence images of U-2 OS cells exposed to hypoxia (1% O_2_, 12 h), showing the localization of 4E-BP1 (red) relative to G3BP1-positive stress granules (SGs; green). Nuclei were counterstained with DAPI (blue). **(B)** Fluorescence intensity line-scan analysis of G3BP1 and 4E-BP1 from (A), showing limited colocalization of 4E-BP1 with hypoxia-induced SGs. **(C)** Representative immunofluorescence images of U-2 OS cells exposed to hypoxia (1% O_2_, 12 h), showing the localization of LARP1 (red) relative to G3BP1-positive SGs (green). Nuclei were counterstained with DAPI (blue). **(D)** Fluorescence intensity line-scan analysis of G3BP1 and LARP1 from (C), showing enrichment of LARP1 within hypoxia-induced SGs. **(E)** Immunoblot analysis confirming siRNA-mediated depletion of LARP1 using two independent siRNAs (siLARP1 #1 and #2) compared with a non-targeting control (siControl). β-actin serves as a loading control. **(F)** Representative immunofluorescence images of U-2 OS cells transfected with siControl or two independent siRNAs targeting LARP1, followed by arsenite treatment (1 h) or hypoxia (1% O_2_, 12 h). SGs were visualized by G3BP1 immunostaining (green), and nuclei were counterstained with DAPI (blue). **(G)** Quantification of SGs per cell from (F), normalized to arsenite-treated controls. **(H)** GFP-Trap co-immunoprecipitation analysis of G3BP1-GFP-expressing U-2 OS cells maintained under unstressed conditions, treated with arsenite (1 h), or exposed to hypoxia (1% O_2_, 12 h). Immunoblotting was performed to detect co-immunoprecipitated LARP1 and CAPRIN1, together with G3BP1-GFP and β-actin. **(I)** Quantification of LARP1 co-immunoprecipitation from (H), normalized to immunoprecipitated G3BP1-GFP and expressed relative to unstressed controls. **(J)** GFP-Trap co-immunoprecipitation analysis of G3BP1-GFP-expressing U-2 OS cells expressing either USP10p^1–40^ or its G3BP1-binding-deficient F10A mutant, USP10p^1–40(F10A)^, under unstressed conditions, arsenite treatment (1 h), or hypoxia (1% O_2_, 12 h). Immunoblotting was performed to assess the association of LARP1 and CAPRIN1 with G3BP1-GFP. **(K)** Quantification of LARP1 co-immunoprecipitation from (J), normalized to immunoprecipitated G3BP1-GFP and expressed relative to unstressed USP10p^1–40^-expressing cells. **(L)** GFP-Trap co-immunoprecipitation analysis of G3BP1-GFP-expressing U-2 OS cells maintained under unstressed conditions, treated with arsenite (1 h), or exposed to hypoxia (1% O_2_, 12 h). Cell lysates were treated with or without RNase A prior to immunoprecipitation to assess the RNA dependence of the G3BP1–LARP1 interaction. CAPRIN1 was included as an additional G3BP1-associated protein. Scale bars, 10 μm (A, C, F). Data are presented as mean ± SD from three independent biological replicates (n = 3). Statistical significance was determined using unpaired Student’s t-tests. ns, not significant; *P < 0.05, ***P < 0.001.

We next considered whether other mTORC1-associated RNA-binding proteins might facilitate SG assembly under hypoxia. One candidate was LARP1, an RNA-binding protein known to interact with the m⁷G cap, 5′ terminal oligopyrimidine (TOP) motifs, and poly(A) tails^43^. LARP1 has also been reported to associate with G3BP1^44,45^ and has been implicated in biomolecular condensation^46^. We therefore examined whether LARP1 localizes to hypoxia-induced SGs. Immunofluorescence analysis revealed robust enrichment of LARP1 within these condensates (Figure 4C–D).

To determine whether LARP1 contributes functionally to SG assembly, we depleted LARP1 and assessed SG formation under hypoxic and arsenite stress conditions. Strikingly, LARP1 depletion strongly impaired hypoxia-induced SG assembly (Figure 4E–G). These findings identify LARP1 as an important regulator of SG formation under hypoxic conditions and suggest that its contribution to condensation may be particularly relevant in this stress context.

Given the established role of G3BP1 as a central SG assembly factor, we next asked whether LARP1 might facilitate SG formation through its association with G3BP1. To investigate this, we performed GFP-Trap immunoprecipitation^47^ using lysates from cells expressing G3BP1-GFP. These experiments confirmed an association between G3BP1 and endogenous LARP1 (Figure 4H–I). Notably, the association was maintained under normoxic and hypoxic conditions but was substantially reduced following arsenite treatment (Figure 4H–I). These observations suggest that the G3BP1–LARP1 association is differentially regulated by stress and may contribute to the distinct assembly requirements of hypoxia-induced SGs.

We next asked whether disruption of G3BP-mediated interactions affects the association between G3BP1 and LARP1. Expression of USP10p^1–40^ significantly reduced G3BP1–LARP1 co-immunoprecipitation during hypoxia (Figure 4J–K), indicating that this association is sensitive to perturbation of G3BP-dependent interactions in a stress-dependent manner. Interestingly, USP10p^1–40^ expression did not substantially disrupt the association between G3BP1 and CAPRIN1, despite previous reports demonstrating competition between these factors^29^. One possible explanation is that the relatively low expression of USP10p^1–40^ achieved through stable expression preferentially disrupts a subset of G3BP-associated interactions while preserving more abundant or higher-affinity complexes.

Finally, because both G3BP1 and LARP1 are RNA-binding proteins, we asked whether RNA contributes to their association. RNase treatment substantially weakened G3BP1–LARP1 co-immunoprecipitation under normoxic, hypoxic, and arsenite conditions (Figure 4L). These findings indicate that RNA contributes to stabilizing the association between G3BP1 and LARP1, potentially through shared RNA interactions or RNA-dependent multivalent complexes.

Together, these results identify LARP1 as a critical regulator of hypoxia-induced SG assembly. The enrichment of LARP1 within hypoxia-induced SGs, its association with G3BP1, and the RNA sensitivity of this interaction support a model in which LARP1 participates in RNA-dependent molecular interactions that facilitate SG formation under hypoxic conditions.

### Hypoxia-induced stress granules exhibit distinct RNA recruitment preferences

Having identified LARP1 as a regulator of hypoxia-induced SG assembly, we next sought to characterize the RNA composition of these condensates and determine whether their recruitment preferences differ from those of canonical arsenite-induced SGs. To accomplish this, we employed Fluorescence-Activated Non-membrane Condensate Isolation (FANCI)^23^ to isolate SGs for transcriptomic analysis.

Briefly, cells expressing G3BP1-GFP were fixed, subjected to mild detergent lysis and sonication, and G3BP1-GFP-positive condensates were isolated by fluorescence-activated sorting. Following isolation, samples were decrosslinked, and RNA was extracted for library preparation and sequencing.

Transcriptomic analysis identified hundreds of RNAs significantly enriched or depleted in hypoxia-induced SG fractions relative to their corresponding input samples (Figure 5A). Similarly, arsenite-induced SGs exhibited extensive enrichment and depletion relative to their corresponding input samples (Figure 5B).

**Figure 5.**
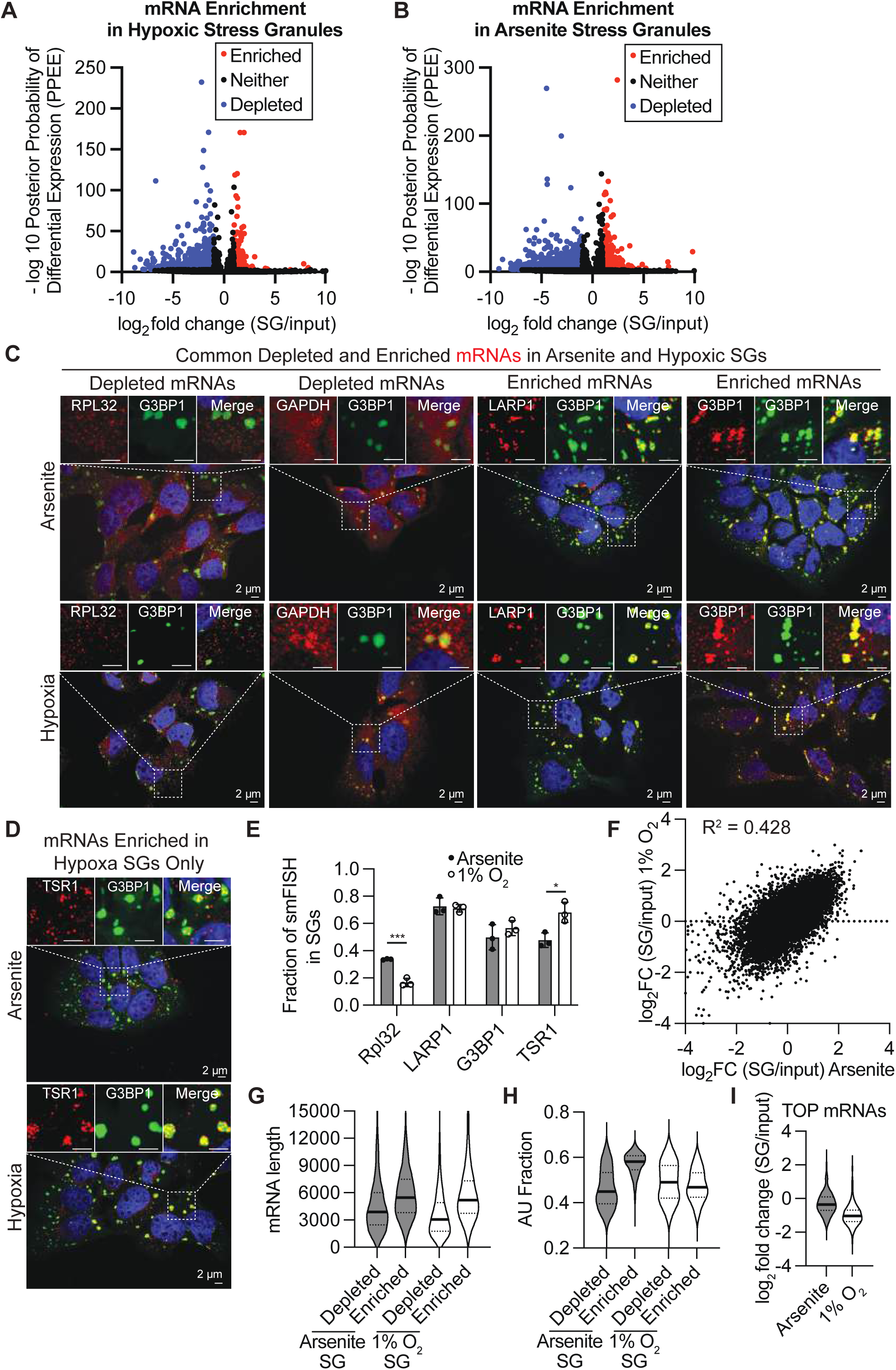
Hypoxia-induced stress granules exhibit conserved RNA recruitment properties but reduced enrichment of TOP mRNAs. **(A, B)** Volcano plots showing the enrichment or depletion of mRNAs in stress granules (SGs) isolated using fluorescence-activated nanoparticle sorting coupled with immunoprecipitation (FANCI) from U-2 OS cells exposed to hypoxia (1% O_2_, 12 h; A) or treated with arsenite (1 h; B). SG-associated RNAs were analyzed by RNA sequencing and compared with their respective input transcriptomes. The x-axis represents log₂ fold change (SG/input), and the y-axis represents −log₁₀ posterior probability of differential expression (PPEE). Enriched, depleted, and non-differentially enriched mRNAs are indicated in red, blue, and black, respectively. **(C)** Representative single-molecule RNA fluorescence in situ hybridization (smFISH) images showing the localization of SG-depleted (RPL32 and GAPDH) and SG-enriched (LARP1 and G3BP1) mRNAs relative to G3BP1-positive SGs in U-2 OS cells treated with arsenite (1 h) or exposed to hypoxia (1% O_2_, 12 h). Target mRNAs are shown in red, G3BP1 immunostaining in green, and DAPI-stained nuclei in blue. Enlarged images of the indicated regions are shown above. Scale bars, 2 μm. **(D)** Representative smFISH images showing the localization of TSR1 mRNA (red) relative to G3BP1-positive SGs (green) in U-2 OS cells treated with arsenite (1 h) or exposed to hypoxia (1% O_2_, 12 h). Nuclei were counterstained with DAPI (blue). Enlarged images of the indicated regions are shown above. Scale bars, 2 μm. **(E)** Quantification of the fraction of smFISH signals localized within SGs for RPL32, LARP1, G3BP1, and TSR1 mRNAs under arsenite (grey) or hypoxia (white) conditions. The fraction was calculated as the number of RNA spots overlapping G3BP1-positive SGs divided by the total number of RNA spots detected per cell. **(F)** Scatter plot comparing transcript enrichment in arsenite-induced and hypoxia-induced SGs, expressed as log_2_ fold change (SG/input). Each point represents an individual transcript. The coefficient of determination (R^2^ = 0.428) indicates a positive correlation in RNA recruitment between the two stress conditions. **(G)** Violin plots showing the transcript length distributions of mRNAs depleted from or enriched in arsenite-induced and hypoxia-induced SGs. **(H)** Violin plots showing the AU nucleotide fraction of mRNAs depleted from or enriched in arsenite-induced and hypoxia-induced SGs. **(I)** Violin plots comparing the enrichment of 5′ terminal oligopyrimidine tract (TOP) mRNAs in arsenite-induced and hypoxia-induced SGs, expressed as log_2_ fold change (SG/input). RNA-seq analyses were performed using three independent biological replicates (n = 3) per stress condition. Data in (E) are presented as mean ± SD from three independent biological replicates (n = 3). Statistical significance in (E) was determined using unpaired Student’s t-tests. *P < 0.05, ***P < 0.001.

To independently validate the transcriptomic measurements, we selected two transcripts classified as SG-enriched and two classified as SG-depleted under both stress conditions and examined their localization by single-molecule fluorescence in situ hybridization (smFISH)^25^. To minimize validation bias, we selected transcripts that had not previously been established as SG-enriched by smFISH^24,48^. Consistent with the sequencing results, transcripts classified as SG-enriched exhibited substantially greater localization to SGs than SG-depleted transcripts under both hypoxic and arsenite stress conditions (Figure 5C,E). We also identified transcripts that exhibited differential SG enrichment between hypoxia and arsenite stress that has not been validated by smFISH. To validate these stress-specific differences, we examined TSR1 mRNA, which was preferentially enriched in hypoxia-induced SGs according to our transcriptomic analysis. Consistent with this prediction, smFISH revealed greater partitioning of TSR1 mRNA into hypoxia-induced SGs than into arsenite-induced SGs (Figure 5D–E). Together, these observations support the reliability of our SG isolation and transcriptomic analyses in identifying both shared and stress-specific RNA recruitment patterns.

Having validated our transcriptomic datasets, we next asked to what extent RNA recruitment into SGs is conserved between hypoxia and arsenite stress. Comparison of SG enrichment scores revealed a positive correlation between the two conditions (R² = 0.428; Figure 5F). These findings indicate that hypoxia- and arsenite-induced SGs share substantial similarities in RNA recruitment, although considerable variation remains between the two stress conditions.

To investigate the molecular features underlying these similarities and differences, we first examined transcript length, a previously established correlate of RNA enrichment in arsenite-SGs^23,24,49–51^. Consistent with earlier studies, longer transcripts were preferentially enriched within hypoxia-induced SGs (Figure 5G). These findings suggest that transcript length is a conserved feature associated with SG recruitment across distinct stress conditions, potentially reflecting increased opportunities for multivalent RNA–RNA and RNA–protein interactions.

We next examined RNA nucleotide composition. Previous studies have reported that RNAs enriched in arsenite-induced SGs exhibit elevated AU content^24,52^. Surprisingly, this relationship was not observed for hypoxia-induced SGs, despite the preferential recruitment of longer transcripts (Figure 5H). Thus, although transcript length remains associated with SG enrichment under hypoxia, the AU-content bias characteristic of arsenite-induced SGs is not conserved. These findings suggest that distinct stress conditions can alter the relative contributions of RNA sequence features to SG recruitment.

Because hypoxia-induced SG assembly depends on the mTORC1–4E-BP1 pathway and requires LARP1, we next asked whether TOP-containing transcripts are preferentially recruited into hypoxia-induced SGs. TOP transcripts encode numerous ribosomal proteins and translation-associated factors^53^ and are well-established targets of mTORC1-dependent translational regulation and LARP1 binding^54^.

Unexpectedly, TOP-containing transcripts were strongly depleted from hypoxia-induced SGs (Figure 5I). This depletion was substantially more pronounced than that observed in arsenite-induced SGs, in which only a subset of TOP transcripts exhibited reduced enrichment. To independently validate this observation, we examined the localization of the TOP-containing transcript RPL32 by smFISH. Consistent with the transcriptomic data, RPL32 exhibited markedly lower SG localization under hypoxia than under arsenite stress (Figure 5C,E).

Collectively, these findings reveal that hypoxia-induced SGs exhibit both conserved and stress-specific RNA recruitment preferences. Although transcript length remains positively associated with SG enrichment, the characteristic AU-content bias of arsenite-induced SGs is diminished under hypoxia. Moreover, the pronounced depletion of TOP-containing transcripts suggests that hypoxia imposes selective constraints on the recruitment of specific RNA classes into SGs.

### Disruption of hypoxia-induced stress granules compromises cell-cycle regulation and cellular fitness

The selective recruitment of RNAs into hypoxia-induced SGs raised the possibility that these condensates regulate the expression of specific functional classes of transcripts. To investigate this, we performed gene ontology analysis of transcripts enriched within hypoxia-induced SGs. This analysis revealed significant enrichment of RNAs associated with rRNA processing, spliceosome function, DNA replication, and cytoplasmic stress granules (Figure 6A). Notably, several transcripts encoding components of the small subunit processome and DNA replication machinery were preferentially enriched within hypoxia-induced SGs.

**Figure 6.**
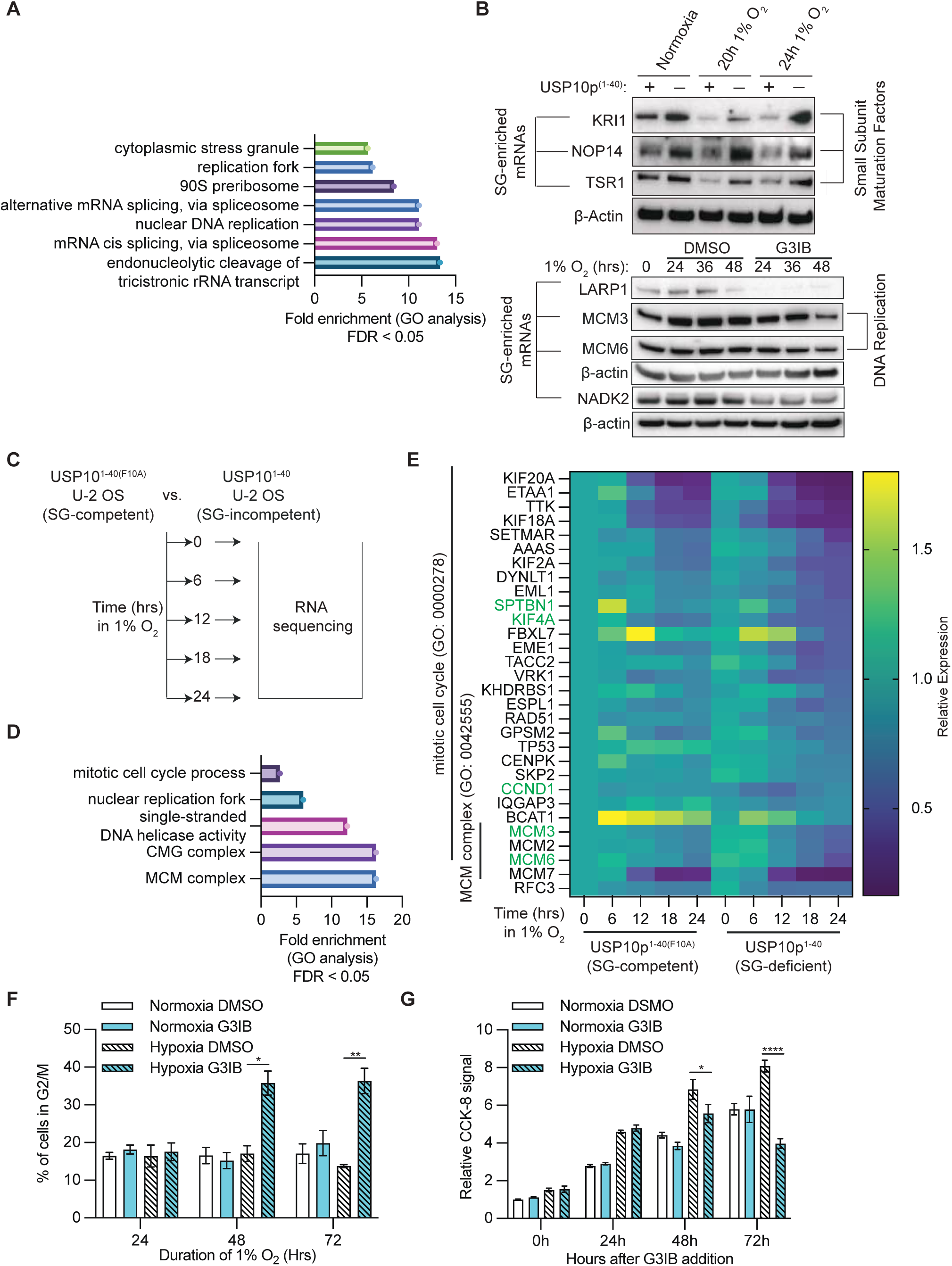
Hypoxia-induced stress granules maintain expression of selected RNAs involved in ribosome biogenesis and DNA replication. **(A)** Gene Ontology (GO) enrichment analysis of transcripts enriched in hypoxia-induced SGs, highlighting biological processes and cellular components associated with RNA processing, ribosome biogenesis, and DNA replication. Bars indicate fold enrichment for significantly enriched GO terms (false discovery rate, FDR < 0.05). **(B)** Immunoblot analysis of proteins encoded by SG-enriched transcripts, including factors involved in small ribosomal subunit maturation (KRI1, NOP14, and TSR1), DNA replication (MCM3 and MCM6), as well as LARP1 and NADK2, in U-2 OS cells subjected to the indicated hypoxic conditions. Upper immunoblots compare cells expressing USP10p^1–40^ or control cells under normoxia and hypoxia (1% O_2_, 20–24 h). Lower immunoblots compare cells treated with DMSO or the SG inhibitor G3IB during prolonged hypoxia (1% O₂, 24–48 h). G3IB was added 10 h after the onset of hypoxia, approximately 2 h before SG assembly. β-actin serves as a loading control. **(C)** Schematic of the experimental design used to investigate SG-dependent RNA regulation during hypoxia. U-2 OS cells expressing either the G3BP1-binding-deficient USP10p^1–40(F10A)^ mutant (SG-competent) or USP10p^1–40^ (SG-deficient) were exposed to hypoxia (1% O_2_) for 0, 6, 12, 18, or 24 h, followed by RNA sequencing to assess changes in transcript abundance over time. **(D)** GO enrichment analysis of transcripts exhibiting significant SG-dependent differential regulation at 12, 18, or 24 h of hypoxia. Differential regulation was determined by comparing transcript abundance at each time point relative to 0 h in SG-competent and SG-deficient cells, followed by calculation of the difference in relative fold changes (ΔΔ fold change; adjusted P < 0.05). Enriched GO terms include mitotic cell-cycle progression, DNA replication, and the minichromosome maintenance (MCM) complex. **(E)** Heatmap showing the relative expression of transcripts associated with mitotic cell-cycle progression (GO:0000278) and the MCM complex (GO:0042555) in SG-competent and SG-deficient U-2 OS cells during hypoxia (0–24 h). Transcript abundance was normalized to the corresponding 0 h condition, and SG-dependent differences are represented as ΔΔ fold changes between SG-competent and SG-deficient cells. Transcripts exhibiting significant differential regulation at 12, 18, or 24 h (adjusted P < 0.05) were included. **(F)** Quantification of the percentage of U-2 OS cells in the G2/M phase of the cell cycle following 24, 48, or 72 h of hypoxia (1% O_2_), in the presence of DMSO or G3IB. Normoxic controls treated with DMSO or G3IB were included for comparison. Cell-cycle distribution was determined by DAPI staining followed by flow cytometry. **(G)** Measurement of cellular metabolic activity using the CCK-8 assay in U-2 OS cells maintained under normoxia or exposed to hypoxia (1% O_2_), in the presence of DMSO or G3IB, over the indicated time course (0–72 h). G3IB was added 10 h after the onset of hypoxia, approximately 2 h before SG assembly. CCK-8 signals were normalized to the 0 h normoxic DMSO-treated control. Data are presented as mean ± SD from three independent biological replicates (n = 3). Statistical significance for the indicated experimental comparisons was determined using unpaired Student’s t-tests. RNA-seq differential regulation was assessed using adjusted P values, with adjusted P < 0.05 considered significant. *P < 0.05, **P < 0.01, ****P < 0.0001.

We next asked whether SG assembly influences the expression of proteins encoded by these transcripts. To address this, we disrupted SG formation using either USP10p^1–40^ or G3IB^21^, a small-molecule inhibitor that targets the same G3BP surface engaged by USP10p^1–40^. Immunoblot analysis revealed that disruption of SG assembly was associated with reduced abundance of several proteins encoded by SG-enriched transcripts, including KRI1, NOP14, TSR1, LARP1, MCM3, MCM6, and NADK2 (Figure 6B). These findings suggest that SG assembly contributes to maintaining the expression of selected proteins under hypoxic conditions.

Given that several of these proteins participate in essential cellular processes, we next investigated whether SG disruption affects broader gene-expression programmes during hypoxia. To address this, we performed RNA sequencing after 12 h of hypoxic exposure and compared transcript abundance between SG-competent and SG-deficient cells (Figure 6C). Gene ontology analysis of differentially expressed transcripts revealed significant enrichment of pathways associated with mitotic cell-cycle processes, DNA replication, and the minichromosome maintenance (MCM) complex (Figure 6D).

Further examination of these gene sets revealed that multiple transcripts involved in cell-cycle regulation and DNA replication exhibited greater reductions in abundance under hypoxia when SG assembly was disrupted by USP10p^1–40^ (Figure 6E). These included several components of the MCM complex, which plays an essential role in DNA replication initiation and replication fork progression^55,56^. Together, these findings suggest that SG-associated regulation of RNA abundance may contribute to maintaining gene-expression programmes required for cell-cycle progression during hypoxic stress.

To determine whether these transcriptome changes were accompanied by alterations in cell-cycle distribution, we examined cells exposed to prolonged hypoxia in the presence or absence of G3IB. Following 48 and 72 h of hypoxic exposure, G3IB-treated cells exhibited an

increased proportion of cells in G2/M compared with control-treated cells (Figure 6F). In contrast, G3IB treatment did not produce a comparable shift in cell-cycle distribution under normoxic conditions. These observations indicate that disruption of SG assembly is associated with altered cell-cycle progression specifically during prolonged hypoxic stress.

We next asked whether these changes were accompanied by reduced cellular fitness. To investigate this, we assessed metabolic activity in cells exposed to hypoxia in the presence or absence of G3IB. G3IB-treated cells exhibited significantly reduced assay signals compared with DMSO-treated cells under hypoxia, whereas this difference was not observed under normoxic conditions (Figure 6H). Notably, the overall assay signal was elevated under hypoxia relative to normoxia, potentially reflecting hypoxia-associated changes in cellular metabolism rather than an increase in viable cell number. Nevertheless, the reduced signal following G3IB treatment suggests that disruption of SG assembly compromises cellular metabolic fitness under hypoxic conditions.

Together, these findings indicate that inhibition of SG assembly is associated with reduced expression of selected proteins involved in RNA processing and DNA replication, altered cell-cycle distribution, and impaired cellular fitness during prolonged hypoxic stress. These observations support a potential role for hypoxia-induced SGs in maintaining gene-expression programmes that contribute to cellular adaptation.

### Hypoxia-induced stress granules protect selected transcripts from RNA decay

Having established that disruption of hypoxia-induced SGs is associated with reduced protein abundance and altered expression of genes involved in essential cellular processes, we next sought to determine how SGs influence gene expression. In particular, we asked whether SGs contribute to maintaining protein abundance by preserving RNA stability, regulating translation, or both.

To address this, we revisited our RNA-sequencing datasets comparing cells expressing USP10p^1–40^ with those expressing the binding-deficient USP10p^1–40(F10A)^ control. We first examined whether transcripts enriched within hypoxia-induced SGs exhibited differential abundance between SG-competent and SG-deficient cells. Interestingly, SG-enriched transcripts on average exhibited higher RNA abundance in cells expressing USP10p^1–40(F10A)^ than in cells expressing USP10p^1–40^ (Figure 7A). In contrast, transcripts classified as SG-depleted did not exhibit a comparable difference in abundance between the two conditions (Figure 7B). These observations suggest that SG recruitment is associated with the maintenance of RNA abundance under hypoxic conditions.

**Figure 7.**
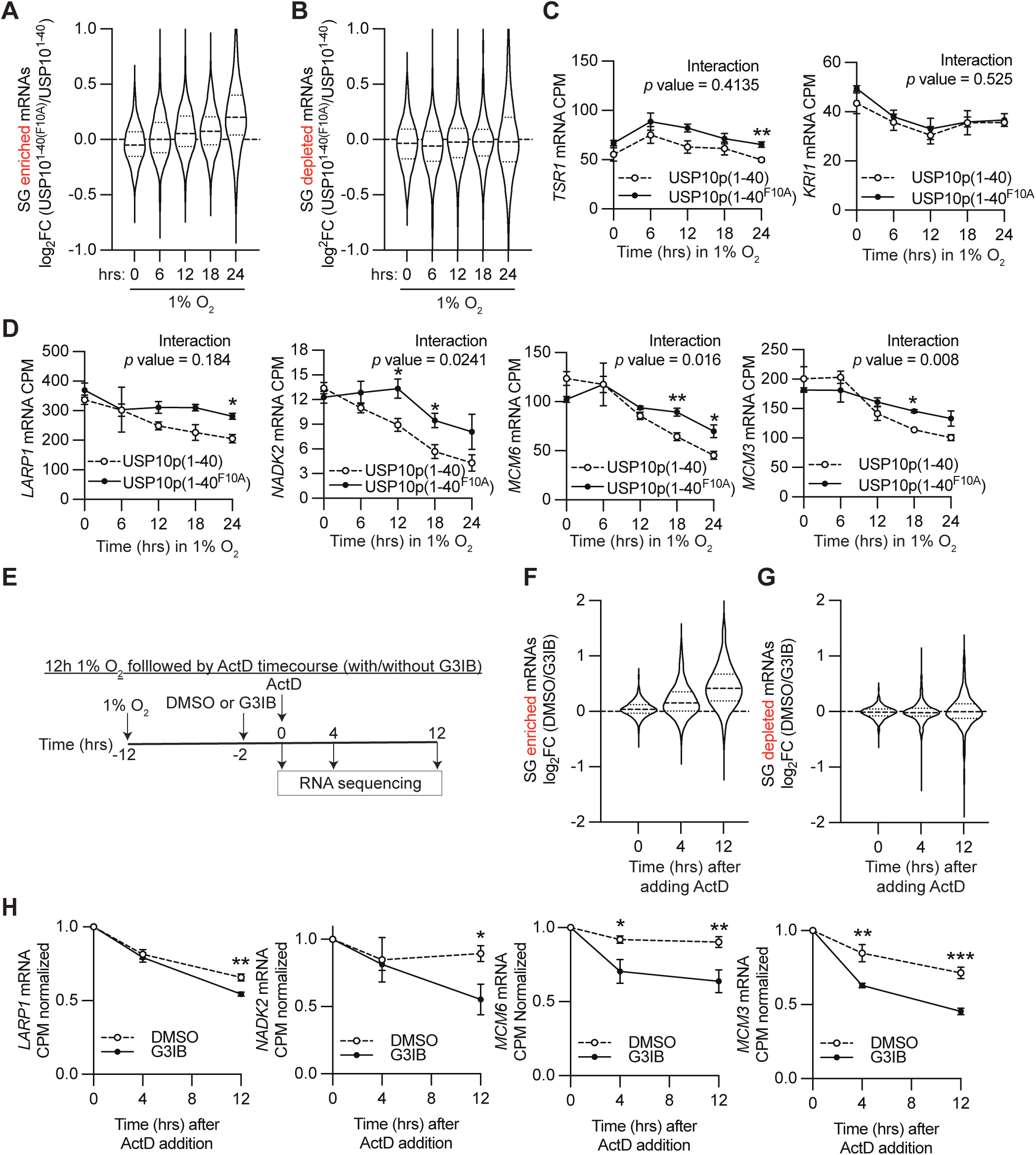
Hypoxia-induced stress granules promote the stability of selected mRNAs during prolonged hypoxia. (A, **B)** Violin plots showing the distribution of log_2_ fold changes in RNA abundance between SG-competent U-2 OS cells expressing USP10p^1–40(F10A)^ and SG-deficient cells expressing USP10p^1–40^ during hypoxia (1% O_2_, 0–24 h). Transcripts were categorized based on their enrichment (A) or depletion (B) in hypoxia-induced SGs, as determined by FANCI-based SG RNA sequencing. Log_2_ fold changes were calculated directly from the ratio of transcript abundance between SG-competent and SG-deficient cells at each time point, without normalization to the corresponding 0 h condition. **(C)** RNA abundance of *TSR1* and *KRI1* in SG-competent U-2 OS cells expressing USP10p^1–40(F10A)^ and SG-deficient cells expressing USP10p^1–40^ during hypoxia (1% O_2_, 0–24 h). Transcript abundance is expressed as counts per million (CPM). Time × cell line interaction P values were determined using a mixed-effects model fitted by restricted maximum likelihood (REML). Statistical comparisons between cell lines at individual time points were performed using unpaired Student’s t-tests. **(D)** RNA abundance of *LARP1*, *NADK2*, *MCM6*, and *MCM3* in SG-competent and SG-deficient U-2 OS cells during hypoxia (1% O_2_, 0–24 h), expressed as CPM. Statistical analyses were performed as described in (C). **(E)** Schematic of the experimental design used to investigate SG-dependent RNA stability during hypoxia. U-2 OS cells were exposed to hypoxia (1% O_2_) for 12 h, with DMSO or the SG inhibitor G3IB added at 10 h, approximately 2 h before SG assembly. Actinomycin D (ActD) was subsequently added at 12 h to inhibit transcription, and RNA was collected at 0, 4, and 12 h following ActD addition for RNA sequencing. **(F, G)** Violin plots showing the distribution of log_2_ fold changes in RNA abundance between DMSO- and G3IB-treated cells at 0, 4, and 12 h following ActD addition. Transcripts were categorized based on their enrichment (F) or depletion (G) in hypoxia-induced SGs. Log_2_ fold changes were calculated directly from the ratio of transcript abundance between DMSO- and G3IB-treated cells at each time point, without normalization to the corresponding 0 h ActD condition. Each transcript is represented individually. **(H)** RNA decay profiles of *LARP1*, *NADK2*, *MCM6*, and *MCM3* following ActD addition in hypoxic U-2 OS cells treated with DMSO or G3IB. Transcript abundance was determined by RNA sequencing and expressed as CPM normalized to the corresponding 0 h ActD condition. Statistical comparisons between DMSO- and G3IB-treated cells at individual time points were performed using unpaired Student’s t-tests. RNA-seq experiments were performed using three independent biological replicates (n = 3). Data in (C, D, H) are presented as mean ± SD. Time × cell line interaction P values in (C, D) were determined using mixed-effects models fitted by REML, whereas individual time-point comparisons in (C, D, H) were performed using unpaired Student’s t-tests. Violin plots in (A, B, F, G) are descriptive, and individual transcripts were not treated as independent biological replicates. *P < 0.05, **P < 0.01, ***P < 0.001.

We next examined whether changes in RNA abundance could account for the reduced protein expression observed following SG disruption. Interestingly, we identified two distinct patterns of regulation. For several transcripts, including KRI1, TSR1, and NOP14, SG disruption reduced protein abundance without a corresponding significant decrease in steady-state RNA levels (Figure 7C). In contrast, other transcripts exhibited reduced RNA abundance that paralleled their decreased protein expression (Figure 7D). These findings suggest that SG-associated regulation of protein expression may involve multiple post-transcriptional mechanisms, including changes in RNA abundance and potentially additional effects on protein production or turnover.

Because differences in steady-state RNA abundance do not necessarily reflect differences in RNA stability, we next directly examined whether SG assembly influences RNA decay. To address this, U-2 OS cells were exposed to hypoxia for 10 h, followed by treatment with G3IB or DMSO. After 12 h of hypoxic exposure, actinomycin D was added to inhibit transcription, and RNA abundance was measured at 0, 4, and 12 h following transcriptional inhibition (Figure 7E).

Analysis of RNA decay revealed that transcripts enriched within hypoxia-induced SGs on average exhibited greater stability in DMSO-treated cells than in G3IB-treated cells (Figure 7F). In contrast, this difference was not observed for SG-depleted transcript population (Figure 7G). These findings suggest that disruption of SG assembly preferentially affects the stability of SG-enriched transcripts rather than causing a uniform increase in RNA decay.

We next examined individual transcripts whose abundance and protein expression were reduced following SG disruption. Notably, LARP1, NADK2, MCM6, and MCM3 transcripts exhibited more rapid decay following G3IB treatment than in DMSO-treated cells (Figure 7H). These results provide direct evidence that pharmacological disruption of SG assembly is associated with reduced stability of selected SG-enriched transcripts under hypoxic conditions. Collectively, these findings support a model in which hypoxia-induced SGs contribute to the maintenance of gene expression by protecting selected transcripts from RNA decay. By preserving RNAs encoding proteins involved in DNA replication, RNA metabolism, and other essential cellular processes, SGs may help sustain the gene-expression programmes required for cellular adaptation to prolonged hypoxic stress.

## Discussion

In this study, we investigated the molecular mechanisms governing SG assembly under hypoxic conditions and their potential roles in cellular adaptation. We demonstrate that hypoxia-induced SGs assemble through a HIF1α –REDD1–mTORC1–4E-BP1 signalling axis, rather than the canonical ISR pathway that drives SG formation under many classical stress conditions. Remarkably, hypoxia-induced SGs assemble despite relatively sustained global protein synthesis, challenging the prevailing association between SG formation and extensive translational repression. We further identify LARP1 as an important regulator of hypoxia-induced SG assembly, suggesting that additional RNA-binding proteins may facilitate condensation under conditions of modest translational inhibition. Transcriptomic characterization revealed that hypoxia-induced SGs share many RNA recruitment preferences with arsenite-induced SGs but exhibit distinct features, including a pronounced depletion of TOP transcripts. Importantly, we provide evidence that SG assembly contributes to the maintenance of selected RNA and protein populations during hypoxia, in part by protecting SG-enriched transcripts from RNA decay. Disruption of SG assembly was further associated with altered cell-cycle distribution and reduced cellular fitness during prolonged hypoxia. Together, these findings support a model in which hypoxia-induced SGs function as selective regulators of post-transcriptional gene expression, potentially contributing to cellular adaptation to sustained oxygen deprivation.

SG formation has traditionally been associated with activation of the ISR^8,30,31^, in which eIF2α phosphorylation suppresses translation initiation and promotes the accumulation of non-translating mRNPs. However, several stress conditions, including selenite exposure^57^, and energy deprivation^58^ and H_2_O_2_^59^, can induce SG assembly through alternative mechanisms involving mTORC1 inhibition and 4E-BP-dependent translational regulation. Our findings extend this framework by demonstrating that hypoxia-induced SG formation depends on the HIF1α –REDD1–mTORC1–4E-BP1 signalling axis and occurs without detectable activation of canonical ISR signalling.

Previous study have implicated both ISR activation and 4E-BP1-dependent mechanisms in hypoxia-induced SG formation, although depletion of individual components of these pathways produced relatively modest effects on hypoxia SG assembly^60^. In contrast, we observed a pronounced dependence on HIF1α, REDD1, and 4E-BP1, whereas pharmacological inhibition of ISR-mediated translational repression had little effect. These differences raise the possibility that the signalling mechanisms governing hypoxia-induced SG assembly vary according to cellular context, oxygen availability, or the duration of hypoxic exposure.

One possibility is that moderate hypoxia preferentially suppresses mTORC1 activity, whereas more severe oxygen deprivation additionally activates the ISR. Under conditions in which both pathways are engaged, it remains unclear whether SG assembly is predominantly governed by one pathway or whether the two mechanisms cooperate to promote condensation. An intriguing possibility is that simultaneous activation of these pathways generates SGs with distinct molecular compositions and regulatory functions. Determining how the relative contributions of ISR and mTORC1 signalling influence SG assembly across different hypoxic conditions may reveal how cells adapt their RNA regulatory responses to varying degrees of oxygen deprivation.

A central principle of SG biology is that translational repression promotes the accumulation of non-translating mRNPs, thereby increasing the availability of RNA substrates capable of undergoing condensation^7^. Consistent with this model, SG formation following arsenite treatment is strongly associated with extensive global translational repression^7,41^. Surprisingly, we found that hypoxia-induced SGs assemble despite relatively sustained global protein synthesis. These findings suggest that extensive translational shutdown is not an absolute requirement for SG formation and that the relationship between translational repression and condensation may depend on the cellular stress context.

One possible explanation is that hypoxia promotes SG assembly through mechanisms that facilitate condensation of a comparatively limited pool of non-translating mRNPs. In this model, SG formation would depend not only on the abundance of untranslated RNAs but also on the molecular interactions that determine their propensity to condense. Such interactions could effectively alter the concentration threshold required for SG assembly, allowing condensation to occur under conditions in which global translation remains largely intact.

Our identification of LARP1 as a regulator of hypoxia-induced SG assembly provides a potential molecular basis for this model. LARP1 is an RNA-binding protein involved in mTORC1-dependent regulation^43^ and has previously been reported to associate with G3BP1^44,45^. We found that LARP1 localizes to hypoxia-induced SGs and is required for their efficient assembly. Moreover, its association with G3BP1 is maintained during hypoxia but reduced following arsenite treatment, suggesting that distinct stress conditions may differentially regulate the molecular interactions contributing to SG condensation. The sensitivity of the G3BP1–LARP1 association to RNase treatment further suggests that RNA contributes to the organization or stability of these complexes.

Although these findings implicate LARP1 in hypoxia-induced SG assembly, its precise molecular function remains unresolved. One possibility is that LARP1 facilitates condensation by increasing the multivalency of RNA–protein interactions within G3BP-containing mRNP complexes. Alternatively, LARP1 may regulate the availability, organization, or recruitment of particular RNA populations that contribute to SG formation. These possibilities are not mutually exclusive and could collectively influence the propensity of non-translating mRNPs to undergo condensation.

Importantly, LARP1 is unlikely to represent the only factor that contributes to SG assembly under conditions of sustained global translation. Additional RNA-binding proteins, changes in cytoplasmic physicochemical properties such as pH^12,61^, or alterations in RNA abundance, processing, modification^48,52^, and structure^62^ may also influence condensation. Defining how these factors interact with G3BP1 and RNA will be important for understanding why hypoxia can induce SG assembly without extensive translational repression. In particular, biochemical reconstitution of LARP1, G3BP1, and RNA may help determine whether LARP1 directly enhances condensation or instead influences SG assembly indirectly through its established roles in RNA regulation.

Our transcriptomic characterization of hypoxia-induced SGs revealed substantial similarities with the RNA composition of arsenite-induced SGs, suggesting that common molecular principles govern RNA recruitment across distinct stress conditions. One conserved feature was the preferential enrichment of longer transcripts, consistent with previous observations that RNA length is an important determinant of SG partitioning^24,49^. Longer RNAs may provide greater opportunities for multivalent RNA–RNA and RNA–protein interactions, thereby increasing their propensity to partition into condensates^63–66^. In particular, the formation of numerous non-specific intermolecular RNA–RNA interactions could contribute to the preferential enrichment of long transcripts independently of specific RNA sequence motifs.

Nevertheless, we also identified important differences between hypoxia- and arsenite-induced SG transcriptomes. Notably, the elevated AU content previously associated with arsenite-induced SG enrichment^24,49^ was not observed under hypoxia. This suggests that although RNA length remains a conserved correlate of SG recruitment, other sequence-dependent or RNA-associated properties may contribute differently across stress conditions. Such differences could arise from changes in RNA-binding protein occupancy, RNA secondary structure, or the translational status of individual transcripts.

One particularly unexpected finding was the pronounced depletion of TOP-containing transcripts from hypoxia-induced SGs. TOP mRNAs encode numerous ribosomal proteins and translation-associated factors and are among the best-characterized targets of mTORC1-dependent translational regulation^38^. Given that hypoxia suppresses mTORC1 signalling and that LARP1 is a major regulator of TOP mRNA translation^54^, the exclusion of these transcripts from hypoxia-induced SGs is especially intriguing.

One possible explanation is that translational repression alone is insufficient to determine whether an RNA partitions into SGs. Although inhibition of translation initiation increases the availability of non-translating mRNPs, their subsequent recruitment may depend on the composition and molecular organization of the associated RNA–protein complexes. In this context, TOP-containing mRNPs may acquire molecular features under hypoxia that favour their retention outside SGs, despite being subject to translational inhibition.

An interesting possibility is that LARP1 participates in distinct RNA regulatory complexes under hypoxia, some of which contribute to SG condensation while others influence the localization or exclusion of particular RNA populations. However, whether LARP1 directly mediates TOP mRNA exclusion remains unknown. Further investigation of the RNA-binding proteins associated with TOP transcripts, together with measurements of their translational status and subcellular localization, may reveal how specific RNA classes are differentially partitioned between SGs and the surrounding cytoplasm.

Collectively, these observations suggest that SG recruitment is governed by a combination of conserved RNA properties and stress-specific molecular interactions. Understanding how these features cooperate to determine RNA partitioning will be important for defining whether SGs formed under different conditions possess distinct regulatory functions.

A longstanding question in SG biology is whether the recruitment of RNAs into SGs alters their subsequent expression or fate^14^. Although SGs have frequently been proposed to function as sites of RNA triage, the extent to which SG localization directly influences RNA stability, translation, or decay remains incompletely understood. Our findings provide evidence that SG assembly under hypoxia is associated with the maintenance of selected RNA and protein populations, supporting a potential role for these condensates in post-transcriptional gene regulation.

We found that several proteins encoded by SG-enriched transcripts were expressed at higher levels in SG-competent cells than in cells in which SG assembly was disrupted.

Interestingly, the underlying changes in RNA abundance differed among transcripts. For some genes, reduced protein abundance was accompanied by lower RNA levels, whereas others exhibited reduced protein expression without a corresponding significant change in steady-state RNA abundance. These findings suggest that SG-associated regulation of gene expression may involve multiple mechanisms, including RNA stabilization and potentially changes in translation or protein turnover.

Consistent with a role in RNA stabilization, our RNA decay experiments revealed that several SG-enriched transcripts, including LARP1, NADK2, MCM3, and MCM6, exhibited greater stability in SG-competent cells than following pharmacological disruption of SG assembly. These observations support the possibility that SGs protect selected transcripts from degradation, thereby contributing to the maintenance of their expression during hypoxic stress. However, whether this protection occurs directly within SGs or results from SG-dependent changes in the composition and activity of associated mRNP complexes remains to be determined.

An intriguing aspect of hypoxia-induced SG biology is the transient nature of their assembly. In our experimental system, SGs emerge after prolonged hypoxic exposure and subsequently disassemble, yet differences in RNA and protein abundance can be observed beyond the period of visible SG formation. This raises the possibility that SGs influence RNA fate not only while transcripts are physically localized within condensates but also after their release into the cytoplasm.

We speculate that SG assembly may promote molecular remodeling of selected mRNPs, generating regulatory states that persist following SG disassembly. Such remodeling could involve changes in RNA-binding protein occupancy, RNA structure, or the association of transcripts with RNA decay and translation machinery. In this model, transient SG localization could establish a form of post-transcriptional memory that influences the subsequent stability or translation of selected RNAs.

This possibility would expand the functional significance of SGs beyond their immediate role as transient sites of RNA condensation. Rather than simply sequestering non-translating mRNPs during stress, SGs could act as dynamic compartments that alter the molecular composition and future regulatory behaviour of enriched transcripts. Determining whether SG-enriched RNAs retain distinct RNA-binding protein interactions or translational properties after SG dissolution will be important for evaluating this model.

Beyond their effects on individual transcripts, our findings suggest that hypoxia-induced SGs may contribute to broader cellular adaptation. Transcriptomic analysis identified SG-associated regulation of genes involved in DNA replication, cell-cycle progression, and ribosomal subunit maturation. Notably, disruption of SG assembly was associated with reduced expression of several MCM complex components and an increased proportion of cells in G2/M during prolonged hypoxic exposure. These changes were accompanied by reduced cellular metabolic fitness, suggesting that SG-associated gene regulation may contribute to maintaining cellular function under sustained oxygen deprivation.

These observations are particularly interesting given the delayed appearance of SGs during hypoxia. Cellular responses to oxygen deprivation occur over multiple temporal phases^67^. Early responses involve rapid changes in metabolism, signalling, and gene expression that enable cells to cope with reduced oxygen availability. In contrast, prolonged hypoxia requires more sustained adaptations that support cellular homeostasis and, in some contexts, continued proliferation despite persistent environmental stress.

The delayed assembly of hypoxia-induced SGs raises the possibility that these condensates participate primarily in this later adaptive phase rather than the immediate response to oxygen deprivation. By regulating transcripts involved in DNA replication, ribosome biogenesis, and other essential cellular processes, SGs may help maintain the gene-expression programmes necessary for cells to tolerate prolonged hypoxic conditions.

One potential connection involves the temporal regulation of HIF-dependent responses. HIF1α and HIF2α can regulate overlapping but distinct transcriptional programmes^68^, with their relative contributions varying according to cell type, oxygen availability, and duration of hypoxia. In certain cellular contexts, HIF2α has been implicated in responses that support prolonged adaptation and proliferation under hypoxic conditions. Our observation suggest SG may intersect with these longer-term HIF-dependent programmes.

We therefore speculate that SGs could provide a post-transcriptional regulatory mechanism that complements HIF-dependent transcriptional responses during prolonged hypoxia. In this model, HIF signalling would contribute to establishing the cellular conditions required for SG assembly, while SGs would help preserve selected transcripts necessary for sustained adaptation. Determining whether SGs preferentially regulate HIF-responsive transcripts or influence the expression of genes associated with prolonged hypoxic adaptation will be an important direction for future investigation.

Hypoxia is a prominent feature of many solid tumours, arising in part from abnormal vascular organization and insufficient oxygen delivery^1^. Hypoxic tumour regions are associated with profound changes in metabolism, gene expression, and cellular behaviour that can contribute to tumour progression and therapeutic resistance. Understanding how cancer cells maintain essential cellular functions under these conditions is therefore important for identifying mechanisms of tumour adaptation and potential therapeutic vulnerabilities.

Our findings suggest that SG assembly may represent an additional layer of post-transcriptional regulation through which cells respond to prolonged hypoxic stress. By preserving selected transcripts involved in DNA replication, RNA metabolism, and other essential cellular processes, hypoxia-induced SGs may help maintain gene-expression programmes that support cellular adaptation to oxygen deprivation. The observation that SG disruption alters cell-cycle distribution and reduces cellular fitness specifically under hypoxic conditions raises the possibility that targeting SG assembly machinery could represent context-dependent vulnerabilities in hypoxic tumour cells. Nevertheless, further studies will be required to establish whether these mechanisms operate broadly across cancer types and contribute directly to tumour growth or survival *in vivo.* In particular, determining whether disruption of hypoxia-induced SG assembly compromises the fitness of tumour cells within hypoxic microenvironments, while sparing cells under normoxic conditions, will be important for evaluating their therapeutic potential.

In summary, our study reveals that hypoxia induces SG assembly through a distinct signalling mechanism that operates despite relatively sustained global translation. We identify LARP1 as an important regulator of this process and demonstrate that hypoxia-induced SGs exhibit general and selective RNA recruitment preferences and are associated with the stabilization of transcripts involved in DNA replication. These findings support a model in which SGs serve not merely as passive repositories of non-translating mRNPs but as dynamic regulators of RNA fate that may contribute to cellular adaptation during prolonged hypoxic stress. Defining the molecular mechanisms connecting SG condensation, RNA remodeling, and sustained gene expression may reveal new principles of post-transcriptional regulation and identify potential vulnerabilities in hypoxic cancers.

## ACKNOWLEDGEMENTS

We thank Dr. Roy Parker and his laboratory (University of Colorado Boulder) for providing the lentiviral plasmids (pRP2985, pRP2986, pRP2987, and pRP2988), and Dr. Paul Taylor and his laboratory (St. Jude Children’s Research Hospital) for providing the G3IB compound. We acknowledge the technical support provided by the GEDAC Core, Spatial Biology Core, and Flow Cytometry Core at the Cancer Science Institute of Singapore (CSI), as well as the Centre for BioImaging Sciences (CBIS) at the National University of Singapore (NUS). We are grateful to Dr. Yilong Zhou for his advice on the FANCI methodology, and to A/Prof Yvonne Tay, Dr. Dennis Kappei, and A/Prof Polly Chen for their valuable discussions and feedback throughout this project. Finally, we acknowledge the financial and institutional support provided by CSI and NUS.

## Notes

### Competing Interest Statement

The authors have declared no competing interest.

